# Molecular and functional characterization of CYP6AN1 orthologs involved in capsaicin metabolism in two *Helicoverpa* species

**DOI:** 10.64898/2026.08.05.743123

**Authors:** Shengyun Li, Zhongyuan Deng, Razak Hussain, Shuang-Lin Dong, Xuguo Zhou, May R. Berenbaum, Xianchun Li

**Author notes:** Corresponding authors: Dr. Xianchun Li; Dr. May R. Berenbaum.

## Abstract

The generalist *Helicoverpa armigera* and the specialist *Helicoverpa assulta* are closely related noctuid pests and are among the few insect herbivores capable of feeding on and damaging hot pepper fruits, which contain the defensive compound capsaicin. Cytochrome P450 monooxygenases (P450s) contribute to the metabolism of plant defensive compounds and can facilitate insect adaptation to chemically defended host plants. Here, we identified *CYP6AN1* in *H. assulta* (*HassCYP6AN1*) and comparatively characterized the *CYP6AN1* orthologs from *H. armigera* (*HarmCYP6AN1*) and *H. assulta*. RACE identified one full-length *HarmCYP6AN1* transcript and three full-length *HassCYP6AN1* transcript isoforms. The *HassCYP6AN1* isoforms contained distinct 5′ UTRs generated by alternative transcription initiation and splicing but shared an identical coding sequence. Sequence comparisons and phylogenetic analysis supported their assignment as orthologs. Constitutive *CYP6AN1* expression was higher in the *H. assulta* midgut, whereas dietary capsaicin significantly induced *CYP6AN1* expression in the *H. armigera* midgut. Recombinant CYP6AN1 proteins were co-expressed with NADPH-cytochrome P450 reductase in *Escherichia coli*, and their substrate-metabolizing activities were evaluated using HPLC-based depletion assays. Both orthologs metabolized capsaicin, but HarmCYP6AN1 exhibited an approximately 2.3-fold higher depletion activity than HassCYP6AN1 under the conditions tested. HarmCYP6AN1 also showed P450-content-dependent xanthotoxin depletion, whereas no detectable xanthotoxin metabolism was observed for HassCYP6AN1. These findings establish CYP6AN1 as a component of the capsaicin-metabolizing repertoire of both species and reveal substantial divergence between the orthologs in transcript organization, expression regulation, catalytic activity, and detectable substrate range.

## 1. Introduction

The cotton bollworm, *Helicoverpa armigera* (Hübner), and the oriental tobacco budworm, *Helicoverpa assulta* (Guenée), are closely related noctuid moths with generally similar morphology (Mitter *et al*., 1993; Fang *et al*., 1997). Both species are widely distributed in Asia, Africa, Australia, and the South Pacific and can cause substantial economic losses (Fitt, 1989; Sharma, 2005; Tay *et al*., 2013). Phylogenetic analyses place the specialist *H. assulta* in an earlier-branching position than the generalist *H. armigera* within *Helicoverpa* (Fang *et al*., 1997; Behere *et al*., 2007; Cho *et al*., 2008). Despite their close relationship and ability to hybridize under laboratory conditions (Wang & Dong, 2001), they differ markedly in host-plant range. *H. armigera* is a highly polyphagous generalist utilizing more than 300 host plant species across 68 families (Cunningham & Zalucki, 2014; Robinson *et al*., 2023), whereas *H. assulta* is an oligophagous specialist restricted primarily to hosts in the Solanaceae, including tobacco (*Nicotiana*) and chili pepper (*Capsicum*) (Chen, 1999; Lee *et al*., 2006; Robinson *et al*., 2023). Both species are important pests of hot pepper and are among the few insects capable of successfully feeding on and damaging pepper fruits (Choi & Boo, 1989; Baek *et al*., 2009; Jia *et al*., 2011).

Chili peppers, which belong to the genus *Capsicum*, are widely cultivated for their distinctive pungency and nutritional value. Their pungency is produced by capsaicinoids, a group of compounds that activate nociceptive receptors in mammals (Caterina *et al*., 1997). Capsaicin is generally the most abundant capsaicinoid and may account for up to 80% of the total capsaicinoid content of pepper fruits (Mazourek *et al*., 2009; Fattori *et al*., 2016). Capsaicinoids can deter some mammalian herbivores and may protect pepper fruits from microbial pathogens and insect herbivores (Thiele *et al*., 2008; Veloso *et al*., 2014; Fattori *et al*., 2016). Capsaicin has been reported to retard larval growth in the spiny bollworm, *Earias insulana* (Weissenberg *et al*., 1986), deter oviposition by the onion fly, *Delia antiqua* (Cowles *et al*., 1989), and inhibit feeding by the phytophagous lady beetle *Henosepilachna vigintioctomaculata* (Hori *et al*., 2011). Natural capsaicinoids also exhibit larvicidal activity against *Anopheles stephensi* and *Culex quinquefasciatus* (Madhumathy *et al*., 2007) and insecticidal activity against *Aphis gossypii* (Li *et al*., 2019). Capsaicin-containing extracts have additionally been reported to enhance the activity of insecticides against *Myzus persicae* and *Leptinotarsa decemlineata* (Edelson *et al*., 2002; Maliszewska & Tęgowska, 2012).

Successful feeding on capsaicinoid-rich pepper fruits by *H. armigera* and *H. assulta* is likely facilitated, at least in part, by metabolic detoxification involving phase I cytochrome P450 monooxygenases (P450s) (Tian *et al*., 2019; Xiong *et al*., 2025) and phase II UDP-glycosyltransferases (UGTs) (Ahn *et al*., 2011). In *H. armigera*, CYP6B6 and three CYP9A enzymes metabolize capsaicin through enzyme-dependent pathways involving hydroxylation, dehydrogenation, and formation of the capsaicin dimer (5, 5′-dicapsaicin (Tian *et al*., 2019). UGT activity mediates capsaicin glucosylation in both *H. armigera* and *H. assulta*, producing capsaicin β-glucoside (Ahn *et al*., 2011). At the tissue level, crude enzyme preparations from the midgut, fat body, and Malpighian tubules—major insect tissues involved in xenobiotic detoxification—metabolize both capsaicin and dihydrocapsaicin in the two species (Zhu *et al*., 2020). More recently, recombinant CYP6B6 from *H. assulta* was shown to metabolize capsaicin and dihydrocapsaicin, providing direct evidence of P450-mediated capsaicinoid metabolism in this specialist species (Xiong *et al*., 2025). Nevertheless, fewer individual capsaicinoid-metabolizing P450s have been functionally characterized in *H. assulta* than in *H. armigera*. Because xanthotoxin, a furanocoumarin occurring in plant taxa within the host range of *H. armigera*, is metabolized by several *H. armigera* P450s, including CYP6AE19, CYP9A12, CYP9A14, and CYP9A17 (Wang *et al*., 2018; Tian *et al*., 2019, Shi *et al*., 2022; Tian *et al*., 2024), it was included as an additional substrate to test whether the *CYP6AN1* orthologs of the generalist *H. armigera* and specialist *H. assulta* differ in detectable substrate range.

In this study, we identified and characterized CYP6AN1 orthologs from *H. armigera* and *H. assulta* to determine their roles in capsaicin metabolism and to assess the extent of molecular and functional divergence between them. We compared their transcript organization, constitutive and capsaicin-induced expression, catalytic activity toward model P450 substrates, and capacity to metabolize capsaicin and xanthotoxin following heterologous co-expression with NADPH–cytochrome P450 reductase. Together, these experiments allowed us to assess whether the two orthologs share a conserved capsaicin-metabolizing function while differing in transcript organization, expression patterns, catalytic activity, and detectable substrate range.

## 2. Materials and methods

### 2.1 Insects and capsaicin induction

Laboratory strains of *H. armigera* and *H. assulta* were established simultaneously from adults that emerged from approximately 2,100 larvae collected from tobacco fields in Xuchang, Henan Province, China, in June 2015. The field-collected larvae, which could not be distinguished reliably by species at the larval stage, comprised approximately 60% *H. armigera* and 40% *H. assulta*. Both species were routinely maintained on wheat germ-based artificial diets (Waldbauer *et al*., 1984) at 27 ± 1 °C under a 16 h light:8 h dark photoperiod. Relative humidity was maintained at 70 ± 10% for adults and 40 ± 10% for larvae.

Analytical-grade capsaicin was purchased from Sigma-Aldrich (USA). Thirty newly molted fifth-instar larvae of each species were fed for 48 h on an artificial diet containing 0.1% (w/w) capsaicin dissolved in methanol. Control larvae were fed the same diet containing an equivalent volume of methanol. Midguts and fat bodies were then dissected separately on ice under a stereomicroscope and washed with ice-cold sterile water. For each species, tissue, and treatment, samples were pooled into three biological replicates of 10 individuals each, flash-frozen in liquid nitrogen, and stored at −80 °C until RNA extraction and analysis of *CYP6AN1* expression.

### 2.2 DNA/RNA extraction and first-strand cDNA synthesis

Genomic DNA (gDNA) was isolated from the midguts of larvae of each species fed the control diet following the procedure described by Sambrook and Russell (2001). Total RNA was isolated from midguts and fat bodies dissected from larvae fed control or capsaicin-containing diets using a guanidinium hydrochloride–based method based on Sambrook et al. (1989). For each biological replicate, 10 pooled midguts or fat bodies were ground to a fine powder in liquid nitrogen and transferred to a 1.5-mL microcentrifuge tube containing 0.6 mL guanidinium hydrochloride RNA extraction buffer. Three biological replicates were prepared for each tissue and treatment. Total RNA was extracted, treated with DNase I (Promega, USA) in the presence of an RNase inhibitor (Thermo, USA) for 30 min to remove residual gDNA, purified by phenol– chloroform extraction and ethanol precipitation, and dissolved in DEPC-treated water. RNA purity was assessed from the A260/A280 and A260/A230 ratios measured using a NanoDrop 2000 spectrophotometer. RNA samples were stored at −80 °C until first-strand cDNA synthesis.

For each sample, 1.2 µg total RNA was reverse-transcribed in a 20-µL reaction containing 2 µL primer mix (Tiangen, China), 4 µL dNTPs, 1 µL M-MuLV reverse transcriptase (New England Biolabs, USA), and 1 µL RNase inhibitor (Thermo, USA). Reverse transcription was performed at 42 °C for 1 h. The resulting first-strand cDNA was stored at −20 °C and subsequently used for PCR cloning and RT-qPCR analysis of *CYP6AN1* expression.

### 2.3 Characterization of CYP6AN1 transcripts by RACE in H. armigera and H. assulta

Partial *CYP6AN1* cDNA sequences from *H. armigera* and *H. assulta* were identified from transcriptomic datasets. Full-length cDNA sequences were obtained by 5′- and 3′-rapid amplification of cDNA ends (RACE) using the SMARTer RACE 5′/3′ Kit (Clontech, USA), following the manufacturer′s instructions with modifications for 3′ RACE. Total RNA was extracted from the midguts of larvae fed control diets.

First-strand 5′- and 3′-RACE-ready cDNAs were synthesized in 20-µL reactions containing 0.5 µg RNA and 200 U SMARTScribe Reverse Transcriptase. The 5′-RACE reaction also contained 1 µL 5′ RACE CDS Primer A and 1 µL SMARTer II A oligonucleotide, whereas the 3′-RACE reaction contained 1 µL 3′ RACE CDS Primer A.

Nested PCR was used to amplify the 5′ and 3′ ends of each *CYP6AN1* cDNA. For 5′ RACE, primary PCR was performed with the Universal Primer A Mix (UPM) and the gene-specific reverse primer R-GSP, followed by secondary PCR with the Universal Primer Short (UPS) and R-NGSP. Each 25-µL reaction contained 12.5 µL 2 × Rapid Taq Master Mix (Vazyme, China), 2.5 µL universal primer, 0.5 µL gene-specific primer (10 µM), and either 1.25 µL 5′-RACE-ready cDNA or 2.5 µL of a 50-fold-diluted primary PCR product. Both rounds of 5′-RACE PCR used a modified touchdown program consisting of 5 cycles at 95 °C for 20 s and 72 °C for 1 min 50 s; 8 cycles each with annealing at 70, 68, and 65 °C for 20 s; and 9 cycles with annealing at 60 °C for 20 s. Each of the latter cycles included denaturation at 95 °C for 20 s and extension at 72 °C for 1 min 30 s, followed by a final extension at 72 °C for 10 min. For 3′ RACE, primary PCR was performed with F-GSP and 3′RP, followed by secondary PCR with F-NGSP and 3′NRP. Each 25-µL reaction contained 12.5 µL 2 × Rapid Taq Master Mix, 1.0 µL each of the forward and reverse primers (10 µM), and either 0.5 µL 3′-RACE-ready cDNA or 1.0 µL of a 50-fold-diluted primary PCR product. Both PCR rounds were conducted for 34 cycles of 95 °C for 15 s, 60 °C for 15 s, and 72 °C for 30 s, followed by a final extension at 72 °C for 10 min. Primer sequences are listed in Table S1.

RACE products were separated on 1.0% agarose gels in 1 × TAE buffer, purified using the AxyPrep DNA Gel Extraction Kit (Axygen, USA), and cloned into the pEASY-T3 Cloning Vector (TransGen Biotech, China). The ligation products were transformed into *Trans1*-T1 competent cells, and plasmids containing inserts of the expected size were sequenced by General Biosystems (Anhui, China).

### 2.4 Cloning of full-length CYP6AN1 cDNA and genomic DNA sequence

The resultant 5′-end and 3′-end sequences, as well as the partial sequence known from the transcriptome data, were then assembled by the DNAMAN software (DNAMAN 6.0.3.99, Canada) into putative full-length *CYP6AN1* transcripts in each species. A pair of primers (M-F/M-R) was designed and used to clone the *CYP6AN1* cDNA of *H. armigera* and three pairs of primers (S-F1/S-R, S-F2/S-R, and S-F3/S-R) were designed and used to clone the *CYP6AN1* cDNA of *H. assulta* based on the 5′- and 3′-RACE results. The PCR reaction mixture contained 12.5 µL 2 × Rapid Taq Master Mix, 1.0 µL forward primer, 1.0 µL reverse primer, and 0.3 µL 3′-RACE-ready cDNA, and the final volume was adjusted with sterile water to 25 µL. The PCR program included an initial denaturation step of 3 min at 95 °C, followed by 35 cycles of 95 °C for 15 s, 50 °C for 15 s, and 72 °C for 45 s, with a final extension of 10 min at 72 °C.

Primers used for cloning full-length *CYP6AN1* cDNA were also employed to clone the *CYP6AN1* genomic sequences of *H. armigera* and *H. assulta* using their gDNA as the template. The total volume of the reaction system was 25 µL, including 12.5 µL of 2 × Rapid Taq Master Mix, 1 µL of each primer (10 µM), and 0.5 µL of gDNA template. The PCR cycling conditions were as follows: 4 min initial denaturation at 94 °C, followed by 35 cycles of 15 s denaturation at 94 °C, 15 s annealing at 50 °C, 2 min 30 s extension at 72 °C, and a 10-min final extension at 72 °C. The PCR products were sequenced at General Biosystems (Anhui, China).

### 2.5 RT-qPCR assays of constitutive and capsaicin-induced expression of CYP6AN1

First-strand cDNA synthesized from midgut and fat-body RNA of *H. armigera* and *H. assulta* larvae was diluted tenfold and used for RT-qPCR analysis of *CYP6AN1* expression. Elongation factor 1 alpha (*EF-1α*) and ribosomal protein L32 (*RPL32*) were used as reference genes. The primers for *EF-1α* and *RPL32* (Table S1) were those described by Zhang et al. (2015). Primer specificity and amplification efficiency were evaluated in accordance with the MIQE guidelines (Bustin et al., 2009).

qPCR was performed using a CFX Connect Real-Time PCR Detection System (Bio-Rad, USA). Each 25-µL reaction contained 12.5 µL 2 × Maxima SYBR Green/ROX qPCR Master Mix (Thermo Scientific, USA), 0.75 µL each of the forward and reverse primers (10 µM; Table S1), 0.8 µL tenfold-diluted cDNA, and 10.2 µL nuclease-free water. The thermal cycling program consisted of an initial denaturation at 95 °C for 10 min, followed by 40 cycles of 95 °C for 15 s, 60 °C for 30 s, and 72 °C for 30 s, with fluorescence acquisition during each cycle. A melting-curve analysis from 65 to 95 °C was performed to verify the amplification of a single product.

A fivefold serial dilution series of cDNA was used to generate a standard curve of Cq values against the logarithm of template concentration for each gene. Amplification efficiency was calculated from the slope of the standard curve as E = 10^−1/slope^− 1 (Bustin *et al*., 2009). Relative *CYP6AN1* expression was calculated using the 2^−ΔΔCq^ method (Livak & Schmittgen, 2001), with normalization to the geometric mean of *EF-1α* and *RPL32* expression values (Vandesompele *et al*., 2002). Three biological replicates were analyzed for each treatment.

### 2.6 Heterologous co-expression of CYP6AN1 orthologs with H. armigera NADPH– cytochrome P450 reductase in E. coli

To obtain functional CYP6AN1 proteins and *H. armigera* NADPH–cytochrome P450 reductase (HarmCPR), their N-terminal sequences were modified before cloning into pCWori for CYP6AN1 or pACYC184 for HarmCPR. Each *CYP6AN1* coding sequence was modified using both the ompA+2 strategy (Pritchard *et al*., 2006) and the 17α strategy (Barnes *et al*., 1991). In the ompA+2 strategy, a cDNA fragment encoding the bacterial ompA leader sequence (MKKTAIAIAVALAGFATVAQA) plus two additional spacer amino acid residues (Ala-Pro) were fused in-frame with the start codons of the two *CYP6AN1s* to preserve their original amino acid sequences (Pritchard *et al*., 2006). In the 17α strategy, the first ten amino acid residues (MIFLLLLPIA) in the N-terminus of each CYP6AN1 were replaced by the N-terminal eight residues (MALLLAVF) of the bovine 17α hydroxylase cytochrome P450 (P45017α) in an attempt to optimize bacterial expression parameters and minimize secondary structure formation in the messenger RNA (Barnes *et al*., 1991). N-terminal modification of HarmCPR was accomplished by in-frame insertion of a cDNA fragment encoding the pelB leader sequence (MKYLLPTAAAGLLLLAAQPAMA) before the start codon of *HarmCPR* cDNA, as described in Pritchard et al. (1997, 2006).

The resultant N-terminus-modified CYP6AN1 and HarmCPR cDNA sequences were cloned into the expression vector pCWori (Kelei Biology, China) via its NdeI and HindIII digestion sites by in-fusion cloning (ClonExpress Ultra One Step Cloning Kit, Vazyme Biotech, China) according to the manufacturer′s instructions. The resulting pCWori-HarmCPR construct was then used as the template to PCR-amplify a DNA segment containing the P_tactac_ promoter of pCWori, *pelB* leader sequence, and *HarmCPR* open reading frame (ORF), which was further ligated into the expression vector pACYC184 (Kelei Biology, China) via its unique BamHI digestion site. The primers used for cloning of the three genes into the two expression vectors are listed in Table S1.

The resultant pACYC184-HarmCPR construct and each of the four pCWori-CYP6AN1 constructs were co-transformed into DH5α competent cells. Single colonies were inoculated into Luria broth (LB) containing ampicillin (50 μg/mL) and chloramphenicol (25 μg/mL) and grown overnight at 37 °C with shaking at 200 rpm. One milliliter of the overnight culture was transferred into 100 mL Terrific broth (TB) supplemented with ampicillin (50 μg/mL), chloramphenicol (25 μg/mL), and thiamine (1 mM), and incubated at 30 °C and 200 rpm. When cultures reached an OD_600_ of 0.7–1.0, ALA (100 μL, 0.5 M) and IPTG (200 μL, 0.5 M) were added to induce protein expression, followed by incubation for 24 h at 30 °C. Cells were harvested by centrifugation at 2,800 g for 20 min at 4 °C. Cell pellets were resuspended in 10 mL ice-cold 1 × TSE buffer [50 mM Tris-acetate (pH 7.6), 250 mM sucrose, 0.25 mM EDTA] containing lysozyme (0.30 mg/mL) and incubated on ice for 45 min. After centrifugation, pellets were resuspended in ice-cold spheroplast resuspension buffer [100 mM potassium phosphate (pH 7.6), 6 mM magnesium acetate, 20% (v/v) glycerol, 0.1 mM DTT, and 1 mM PMSF], sonicated on ice, and centrifuged at 14,000 g for 12 min at 4 °C. The supernatant was ultracentrifuged at 180,000 g for 1 h at 4 °C, and the membrane fraction was resuspended in 1 × TSE buffer and stored at −80 °C for subsequent enzyme assays.

Total protein concentration of each CYP6AN1/HarmCPR enzyme source was determined according to Bradford (1976). The HarmCPR activity was assayed by its NADPH–cytochrome *c* reduction activity according to the method described in Pritchard *et al*. (2006) using an extinction coefficient of 21.4 mM^−1^ cm^−1^ at 550 nm. Reduced CO-difference spectra assays (Omura & Sato, 1964; Guengerich *et al*., 2009) were performed to determine if the expressed CYP6AN1 of each CYP6AN1/HarmCPR enzyme preparation was functional (yielding a peak at 450 nm) and to measure its content using an extinction coefficient of 0.091 μM^−1^ cm^−1^. The CYP6AN1/HarmCPR enzyme preparations with functional CYP6AN1 were analyzed to reveal their ability to metabolize two P450 model substrates, 7-ethoxycoumarin and 7-methoxyresorufin, as well as two phytochemicals, capsaicin and xanthotoxin.

### 2.7 Enzymatic activities of heterologously co-expressed CYP6AN1/HarmCPR enzymes towards two P450 model substrates

The 7-ethoxycoumarin O-deethylation (ECOD) and 7-methoxyresorufin O-demethylation (MROD) activities of heterologously co-expressed CYP6AN1/HarmCPR were analyzed using the spectrofluorometric method described by Waxman and Chang (2006). All assays were performed in a 2.0-mL microtube. The assay mixture for ECOD contained 186 μL of ECOD assay buffer [100 mM potassium phosphate (pH 7.4), 20% (v/v) glycerol, and 0.1 mM EDTA], 1 μL of 7-ethoxycoumarin (200 mM), and 7 μL of membrane fraction. The reaction was preincubated to 37 °C and initiated by adding 5.0 μL Solution A and 1.0 μL Solution B (NADPH Regeneration System; Promega, USA) in a shaker at 200 rpm for 20 min. The reaction was stopped on ice and 25 μL of 2 M hydrochloric acid was added, after which 230 μL of chloroform was added. The reaction solution was centrifuged for 5 min at 3,000 g. The upper aqueous layer was removed, and the organic layer (200 μL) was transferred to a new 1.5-mL tube. Then 125 μL of sodium borate (30 mM) was added to the tube and the reaction mixture was centrifuged for another 5 min at 3,000 g. A 100-µL aliquot of the upper layer was used to measure the enzyme activities at an excitation wavelength of 370 nm (slit 5 nm) and an emission wavelength of 450 nm (slit 5 nm).

A negative control with corresponding preparation from cells carrying the empty vectors pCWori and pACYC184 was set up in parallel. The reactions with the heat-inactivated membrane fraction (inactivated by incubation at 95 °C for 5 min)/without NADPH addition were also used as controls. Each assay was performed in three independent reactions. Tubes containing a known amount of serially diluted metabolite (7-hydroxycoumarin) and assay buffer were subjected to the same treatments. A standard curve was generated to quantify the amount of 7-hydroxycoumarin in the product. Velocities were calculated based on fluorescence values within linear range and presented as the formation of 7-hydroxycoumarin. The final metabolic activity was corrected by subtracting the background (negative control) and normalized to P450 content. Accordingly, for determination of P450-catalyzed MROD activities, a reaction mixture containing MROD assay buffer [100 mM potassium phosphate (pH 7.4), 1.5 mM EDTA] was used to measure the fluorescence at an excitation wavelength of 530 nm (slit 5 nm) and an emission wavelength of 582 nm (slit 5 nm). Authentic metabolite (resorufin) was used as standards for calculating activity of each unknown sample.

### 2.8 In vitro metabolism of capsaicin and xanthotoxin by heterologously co-expressed CYP6AN1/HarmCPR enzyme preparations

Separate methanolic stock solutions of capsaicin and xanthotoxin (Sigma-Aldrich, USA) were prepared. Before use, the capsaicin stock was diluted tenfold with 0.1 M sodium phosphate buffer (pH 7.4), yielding a 0.7 mM working solution; the xanthotoxin stock concentration was 0.8 mM. The reaction mixture, in a final volume of 200 μL, consisted of 0.1 M sodium phosphate buffer (pH 7.4), 5.0 μL Solution A and 1.0 μL Solution B (NADPH Regeneration System; Promega, USA), substrate (10 nmol capsaicin or 1.43 nmol xanthotoxin), and a membrane fraction containing recombinant CYP6AN1/HarmCPR. Four P450 amounts were tested for each enzyme–substrate combination to evaluate substrate depletion across a range of P450 contents. For capsaicin, HarmCYP6AN1 was tested at 2.57, 5.13, 10.25, and 20.50 pmol P450 per reaction, whereas HassCYP6AN1 was tested at 4.90, 9.80, 14.71, and 19.61 pmol P450 per reaction. For xanthotoxin, HarmCYP6AN1 was tested at 2.57, 7.70, 10.25, and 15.37 pmol P450 per reaction, whereas HassCYP6AN1 was tested at 2.45, 4.90, 9.80, and 19.61 pmol P450 per reaction.

Reaction mixtures were preincubated in a water bath at 30 °C for 5 min, and reactions were initiated by the addition of substrate. The reactions were then incubated on an orbital shaker at 30 °C and 250 rpm for 30 min, a duration selected based on preliminary time-course experiments showing progressive substrate depletion over this interval. Reactions were terminated by adding 200 µL ice-cold acetonitrile, followed by shaking at 30 °C and 250 rpm for 5 min. Samples were centrifuged at 12,000 × g for 5 min and filtered through 0.22-µm filters into amber vials for immediate HPLC analysis. For inhibition assays, commercial liquid piperonyl butoxide (PBO; Bide Pharmatech, China) was diluted tenfold with 0.1 M sodium phosphate buffer (pH 7.4) immediately before use and the freshly prepared dispersion was vortexed thoroughly before pipetting. A 5.2-µL aliquot of the tenfold-diluted PBO dispersion was added to each 200-µL reaction, corresponding to a nominal final concentration of approximately 8 mM. Membrane fractions were mixed with PBO before being added to the reaction mixture.

Reactions containing equivalent amounts of empty-vector membrane fractions prepared from *E. coli* cells carrying pCWori and pACYC184 were conducted concurrently to assess background substrate loss unrelated to recombinant CYP6AN1 activity. Each enzyme–substrate combination was tested in triplicate at each P450 amount. Residual capsaicin and xanthotoxin were analyzed using an HPLC system equipped with a UV detector (Waters Corporation, USA) and a reversed-phase C_18_ column (Agilent, USA; 4.6 × 250 mm, 5-µm particle size). For capsaicin analysis, the mobile phase consisted of methanol and water (71:29, v/v), modified from the methods of Juangsamoot *et al*. (2012) and Henderson *et al*. (1999), and capsaicin was detected at 280 nm. For xanthotoxin analysis, the mobile phase consisted of methanol and water (50:50, v/v), and xanthotoxin was detected at 297 nm. For both substrates, the flow rate was 0.8 mL/min, the injection volume was 15 µL, and the column temperature was maintained at 30 °C.

Calibration standards containing known amounts of capsaicin or xanthotoxin were prepared in parallel with the enzyme-reaction samples and subjected to the same sample-processing procedures. Linear calibration curves relating HPLC peak area to the known amount of each substrate were generated, and the amount of substrate remaining in each reaction was determined from the corresponding calibration curve. For each CYP6AN1 preparation and P450 content, the mean substrate amount measured in the three PBO-treated reactions was used as the inhibited control. Substrate depletion in each corresponding reaction without PBO was calculated by subtracting the amount remaining in that reaction from the mean amount measured in the PBO-treated controls. Specific substrate-depletion activity was calculated by normalizing substrate depletion to incubation time and P450 content and was expressed as pmol substrate/min/pmol P450. Measurements obtained at the highest P450 content were excluded because substrate depletion no longer increased proportionally with P450 content, indicating that these measurements were outside the proportional enzyme-content range of the assay.

### 2.9 Phylogenetic analysis of CYP6AN proteins

Amino acid sequences of 19 CYP6 proteins from *H. armigera*, *H. assulta*, *Helicoverpa zea*, *Spodoptera frugiperda*, *S. littoralis*, *S. litura*, *S. exigua*, *Chrysodeixis includens*, and *Trichoplusia ni* were included in the phylogenetic analysis. The dataset contained the focal HarmCYP6AN1 and HassCYP6AN1 proteins, closely related CYP6AN paralogs, representative CYP6AN-like proteins from other noctuid species, and *H. armigera* CYP6AE14 and *H. armigera* CYP6AE16 as outgroup sequences. Because relatively few annotated *H. assulta* CYP6 sequences were available in public protein databases, the *H. assulta* genome assembly ASM2961881v1 (GCA_029618815.1) was searched using identified HassCYP6AN1 protein sequences as queries. Candidate genomic regions were identified based on BLAST similarity and subsequently examined through gene prediction, sequence alignment, and genomic organization. The sequences are provided in Supplementary file 1.

Protein sequences were aligned using MUSCLE implemented in MEGA 11 (Molecular Evolutionary Genetics Analysis version 11; Tamura *et al*., 2021) with the default amino-acid alignment parameters. The optimal amino-acid substitution model was determined by maximum-likelihood model testing in MEGA 11 according to the Bayesian information criterion. The LG model with gamma-distributed rate variation among sites—LG+G—provided the best fit to the data (BIC = 12,991.830; AICc = 12,735.268; lnL = −6,331.490). A maximum-likelihood phylogeny was subsequently reconstructed using the LG+G model. Rate variation among sites was approximated using five discrete gamma categories. Positions containing gaps or missing data were excluded using complete deletion. The initial tree was generated automatically using the neighbor-joining/BioNJ method, and tree topology was optimized using nearest-neighbor interchange. Branch support was evaluated using 1,000 bootstrap replicates. The final tree was displayed as a phylogram, with branch lengths representing the estimated number of amino-acid substitutions per site.

### 2.10 Sequence and statistical analysis

Amino acid sequences of the two CYP6AN1 proteins were compared with bacterial P450 101A (P450_cam_) and rat P450 2A1 (Gotoh, 1992) to map the six substrate recognition sites (SRSs). The transcript sequences of the two *CYP6AN1* orthologs obtained by RACE were aligned with their gDNA sequences obtained by PCR to locate the exon and intron regions and their boundaries in two species using NCBI BLAST-Global Align. The sequences used for this analysis are provided in Supplementary file 2.

Significant differences in the expression levels of *CYP6AN1* orthologs in the midgut and fat body of the two species were determined by one-way analysis of variance (ANOVA) followed by Tukey′s HSD tests for multiple comparisons. Two-tailed Student′s *t*-tests were used to compare *CYP6AN1* expression levels between methanol-control and capsaicin-treated larvae within each species and tissue. Two-tailed Student′s *t*-tests were also used to compare the enzymatic activities of HarmCYP6AN1 and HassCYP6AN1 toward each of the two model substrates and capsaicin.

## 3. Results

### 3.1 Characterization of CYP6AN1 transcripts and encoded proteins in H. armigera and H. assulta

RACE analysis identified multiple 5′-end and full-length *CYP6AN1* transcript isoforms in *H. assulta* (Fig. 1A and B), whereas only a single full-length transcript isoform was detected in *H. armigera* (Fig. 1B). The full-length *H. armigera CYP6AN1* transcript was 2,285 bp in length and contained a 5′ UTR of 409 bp, a 3′ UTR of 319 bp, and an open reading frame (ORF) of 1,557 bp encoding a deduced protein of 518 amino acids (aa) (Fig. 2). Alignment of the full-length transcript with its genomic DNA sequence revealed that the gene contains two introns located in the 5′ UTR and ORF, respectively, both of which conform to the canonical GT/AG splicing rule (Fig. 1B). Protein BLAST (BLASTP) analysis using the deduced amino acid sequence of this transcript as the query identified the protein (GenBank protein accession no. AID54897.1) encoded by the previously reported *CYP6AN1* transcript from *H. armigera* populations in Pakistan and Australia (GenBank accession no. KM016745.1; Rasool *et al*., 2014) as the highest-scoring match. The two proteins shared 98.6% amino acid identity. Based on the high sequence similarity and its identification as the best BLASTP hit to *H. armigera* CYP6AN1, the transcript was assigned the name *CYP6AN1v2* (GenBank accession no. PQ470130.1) by David Nelson (personal communication) and is hereafter referred to as *HarmCYP6AN1*.

**Fig. 1.**
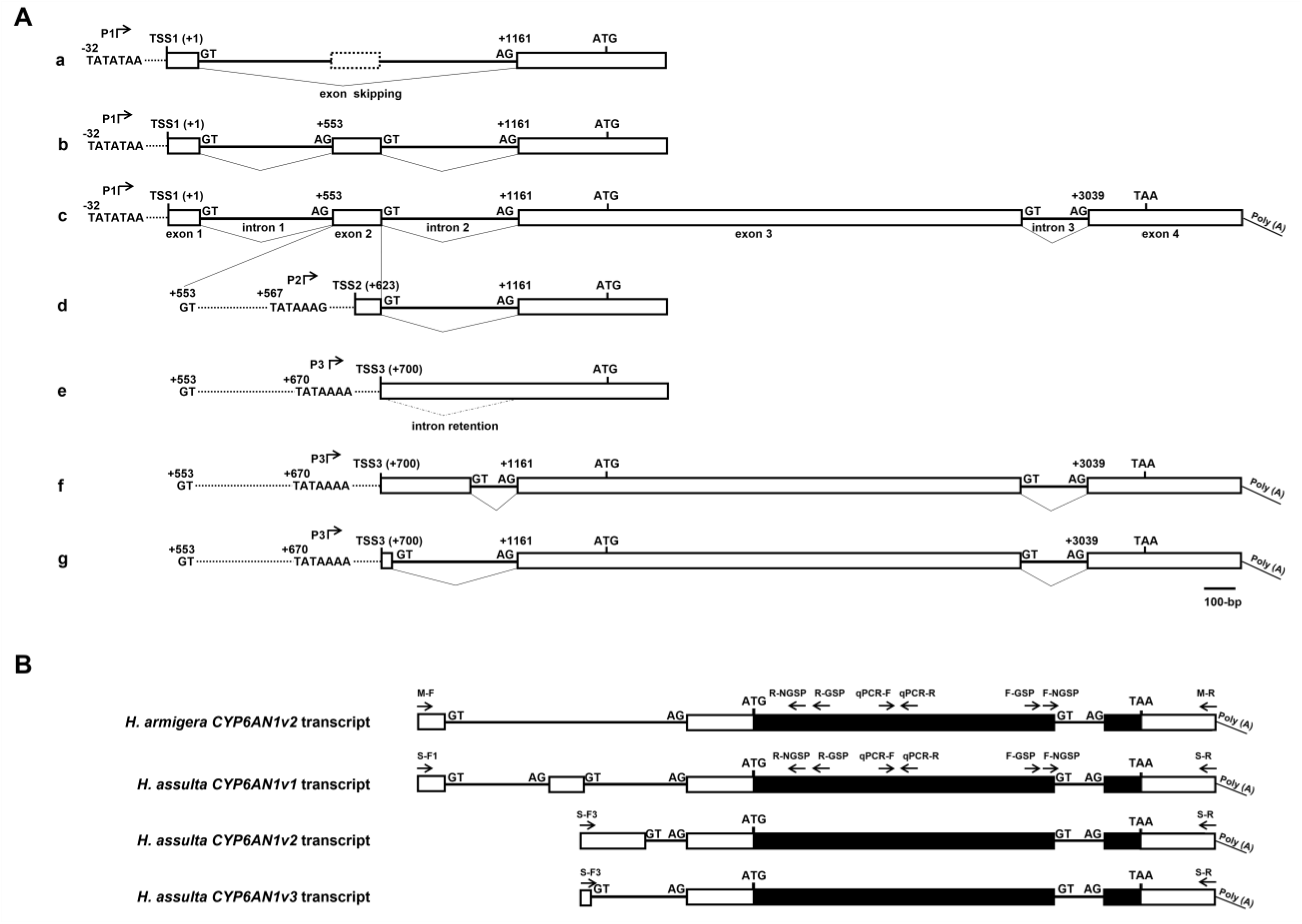
Diagram of 5′-RACE end and full-length cDNA sequences of *H. assulta CYP6AN1* and *CYP6AN1* transcripts in *H. armigera* and *H. assulta*. A: Diagrams a, b, d, and e represent 5′-RACE sequences. Diagram c, f, and g represent full-length cDNA sequences. P1, P2, and P3 represent three alternative promoter regions which contain TATA-box elements. White boxes and bold solid lines between boxes show the exons and introns of the gene, respectively. Splice sites GT/AG, start codon ATG, stop codon TAA, and Poly(A) tail are also indicated in the figure. The start position of each TATA-box and exon is numbered relative to the transcription start site 1 (TSS1, indicated by +1) of the sequence (a, b, and c), with sequence upstream of it preceded by “-”, and downstream of it preceded by “+”. Dotted lines represent omitted nucleotides and were not shown to scale. B: Boxes and lines between boxes represent exons and introns of genes, respectively. White and black boxes represent untranslated regions (UTRs) and protein-coding regions of the exons, respectively. The annealing directions and relative positions of the primers used for the 5′/3′ RACE, full-length cDNA cloning, and RT-qPCR analyses of gene expression are depicted with arrows and the corresponding primer names. GT/AG indicates the canonical splice sites. Start codon ATG, stop codon TAA, and Poly (A) tail are also indicated (the diagrams are schematic and not drawn to scale).

**Fig. 2.**
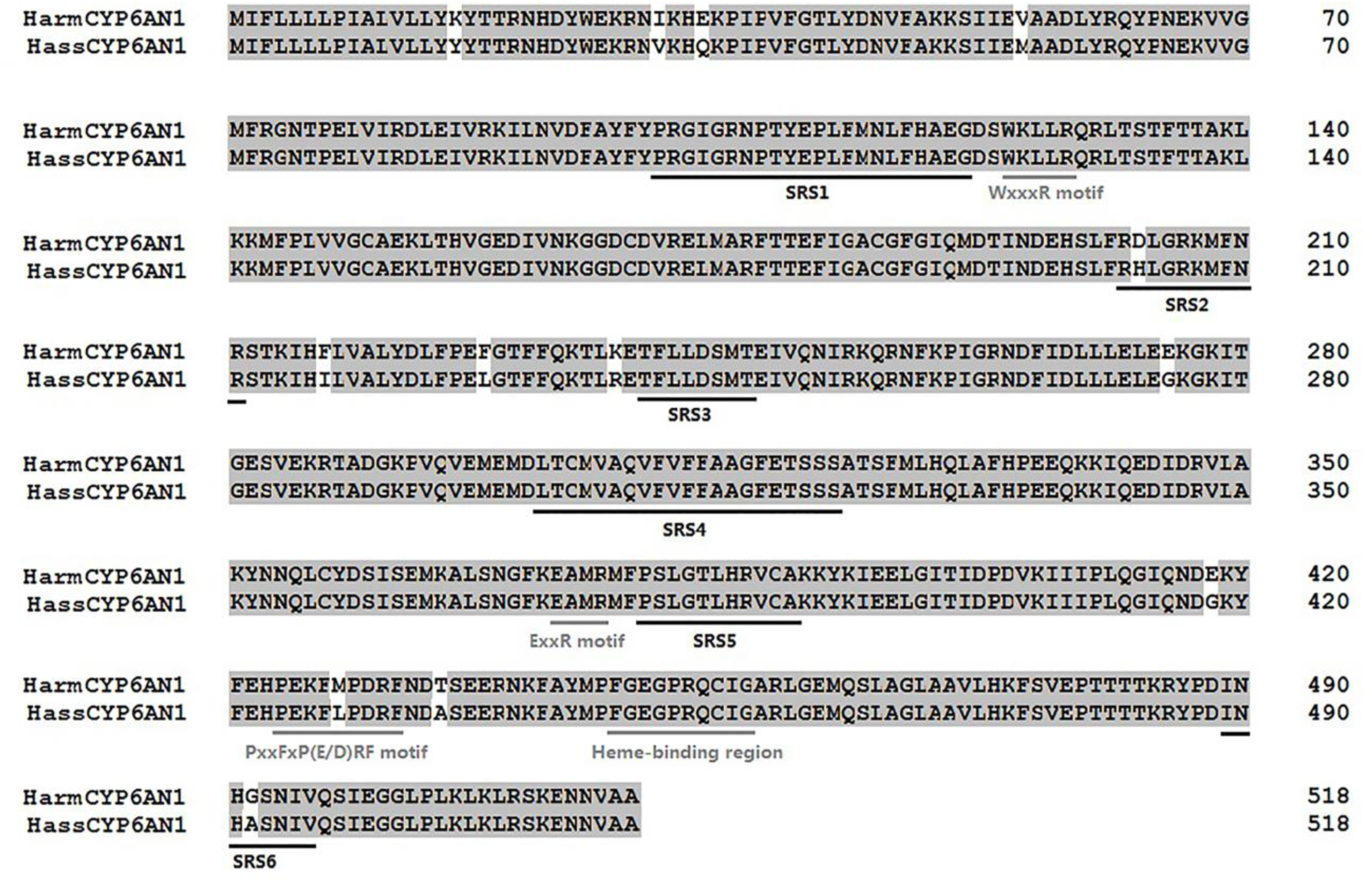
Amino acid sequence alignment between HarmCYP6AN1 and HassCYP6AN1. Black underlines indicate putative substrate recognition sites (SRSs). Gray underlines indicate consensus sequences of the conserved P450 motifs: helix-C (WxxxR), helix-K (ExxR), PxxFxP(E/D)RF motif, and the heme-binding region (FxxGxRxCxG), where x is any amino acid.

The *H. assulta CYP6AN1* gene (hereafter referred to as *HassCYP6AN1*) was transcribed into at least 6 different transcripts (b and c in Fig. 1A represented the same transcript); each contained a 5′ UTR of different length but a common 333-bp 3′ UTR and a common 1,557-bp ORF encoding a deduced protein of 518 amino acids (Fig. 2). Alignment of the RACE products with the genomic sequence suggested that alternative transcription initiation near three putative TATA-box-associated promoter regions (named P1, P2, and P3; Fig. 1A and S1), together with alternative splicing within the 5′ UTR, generated the observed transcript structures. The 5′-RACE products a and b and full-length transcript c were inferred to initiate at TSS1 (+1), adjacent to the 5′-most putative promoter region, P1. The 5′-RACE product d initiated at TSS2 (+623 relative to TSS1), adjacent to the putative P2 region, resulting in a 375-bp 5′ UTR. Relative to 5′-RACE product b and full-length transcript c, 5′-RACE product a lacked exon 2 because of exon skipping, producing a 395-bp 5′ UTR. The 5′-RACE product e and full-length transcripts f and g were inferred to initiate at TSS3 (+700), adjacent to the putative P3 region (Fig. 1A). These three transcripts were generated by alternative 5′ splice-site selection within intron 2 (f vs. g) or retention of intron 2 (e), yielding a 5′ UTR of 751 bp (e), 595 bp (f), and 336 bp (g), respectively (Fig. 1A). These three full-length *HassCYP6AN1* transcripts were designated *CYP6AN1v1* (GenBank accession no. PQ470131.1), *CYP6AN1v2* (GenBank accession no. PQ470132.1), and *CYP6AN1v3* (GenBank accession no. PQ470133.1), respectively (David Nelson, personal communication). All three full-length transcripts encode the same 518-aa protein. BLASTP analysis using HassCYP6AN1 as the query identified *H. armigera* CYP6AN1 (GenBank protein accession no. AID54897.1), encoded by the previously reported transcript KM016745.1, as the highest-scoring match. These findings support the assignment of *HarmCYP6AN1* as the putative *H. armigera* ortholog of *HassCYP6AN1*. Both proteins contain the conserved P450 heme-binding motif FxxGxRxCxG, C-helix WxxxR motif, K-helix ExxR motif, and post-K-helix PxxFxP(E/D)RF motif. Of the 13-aa substitutions, two occur within predicted substrate recognition sites (SRSs): H203D in SRS2 and A492G in SRS6 (Fig. 2).

### 3.2 Phylogenetic relationships of CYP6AN proteins

Maximum-likelihood analysis resolved a CYP6AN-related clade that included several proteins provisionally annotated in NCBI as CYP6B5-like or CYP6B6-like. This clade was distinct from the CYP6B2-like group and the CYP6AE outgroup sequences (Fig. 3). The CYP6AN-like proteins from *Helicoverpa*, *Spodoptera*, *Chrysodeixis*, and *Trichoplusia* formed a strongly supported clade with 100% bootstrap support. Within *Helicoverpa*, HassCYP6AN1 clustered with HarmCYP6AN1 and a *H. zea* P450 6B6-like protein (XP_047019984.1), with 100% bootstrap support for the three-sequence clade (Fig. 3). HarmCYP6AN1 and *H. zea* P450 6B6-like formed a sister pair with 60% bootstrap support.

**Fig. 3.**
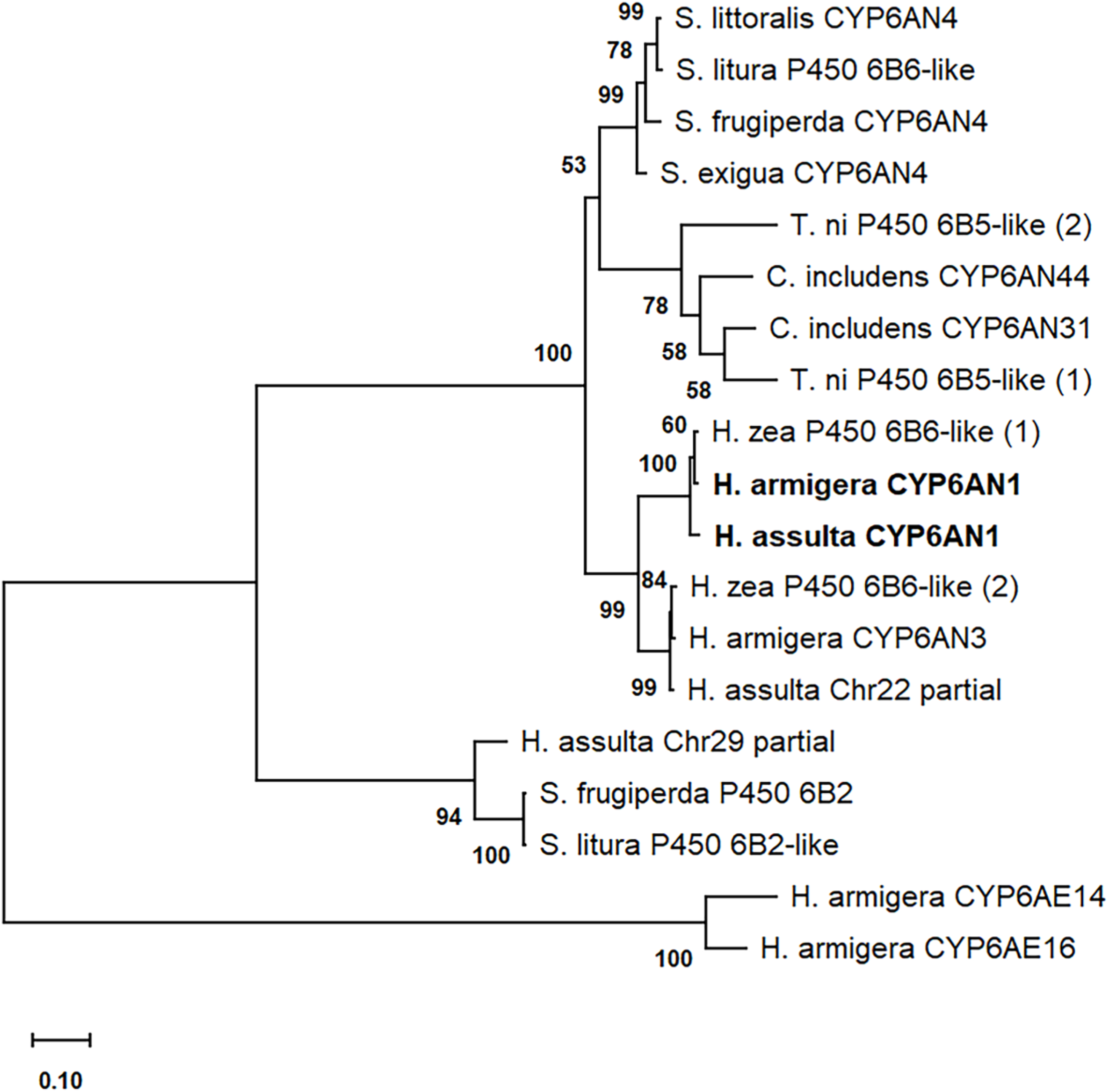
Maximum-likelihood phylogeny of selected noctuid CYP6 proteins. Amino acid sequences were aligned using MUSCLE, and the phylogeny was reconstructed in MEGA 11 using the LG model with gamma-distributed rate variation among sites. Positions containing gaps or missing data were excluded by complete deletion. Bootstrap support values from 1,000 replicates are shown at the internal nodes. *H. armigera* CYP6AE14 and CYP6AE16 were used as outgroup sequences. Branch lengths represent the estimated number of amino acid substitutions per site. Numbers in parentheses distinguish proteins sharing the same provisional annotation in the NCBI database. The focal *H. armigera* CYP6AN1 and *H. assulta* CYP6AN1 proteins are shown in bold. “Chr22” and “Chr29” denote protein sequences derived from chromosomes 22 and 29, respectively, of the *H. assulta* genome assembly ASM2961881v1.

The strongly supported grouping of the CYP6AN1-like proteins from *H. assulta*, *H. armigera*, and *H. zea* supports their assignment to the same orthologous lineage. A distinct *Helicoverpa* lineage contained the partial *H. assulta* chromosome 22 sequence, *H. armigera* CYP6AN3, and another *H. zea* P450 6B6-like protein (XP_047019985.1).

This three-sequence clade received 99% bootstrap support and was clearly separated from, but closely related to, the CYP6AN1 orthologous group, suggesting that it represents a paralogous lineage derived from an ancestral gene duplication. *H. armigera* CYP6AE14 and CYP6AE16 formed a strongly supported outgroup pair with 100% bootstrap support. Overall, the phylogenetic topology strongly supports the assignment of HassCYP6AN1 and HarmCYP6AN1 to the same CYP6AN1 orthologous lineage and clearly distinguishes this lineage from the separate paralogous lineage.

### 3.3 Constitutive and capsaicin-induced expression of CYP6AN1 in H. armigera and H. assulta

RT–qPCR analysis showed that the constitutive expression of *CYP6AN1* in larval midgut and fat body was significantly higher in the specialist *H. assulta* than in the generalist *H. armigera*, with a 2.7-fold increase in the midgut and a marginal 1.5-fold increase in the fat body (Fig. 4A). Capsaicin treatment significantly induced *CYP6AN1* expression in the midgut of fifth-instar *H. armigera* larvae (4.5-fold; Fig. 4B), whereas no significant induction was observed in the fat body of *H. armigera* or in either tissue of *H. assulta* (Fig. 4B and C).

**Fig. 4.**
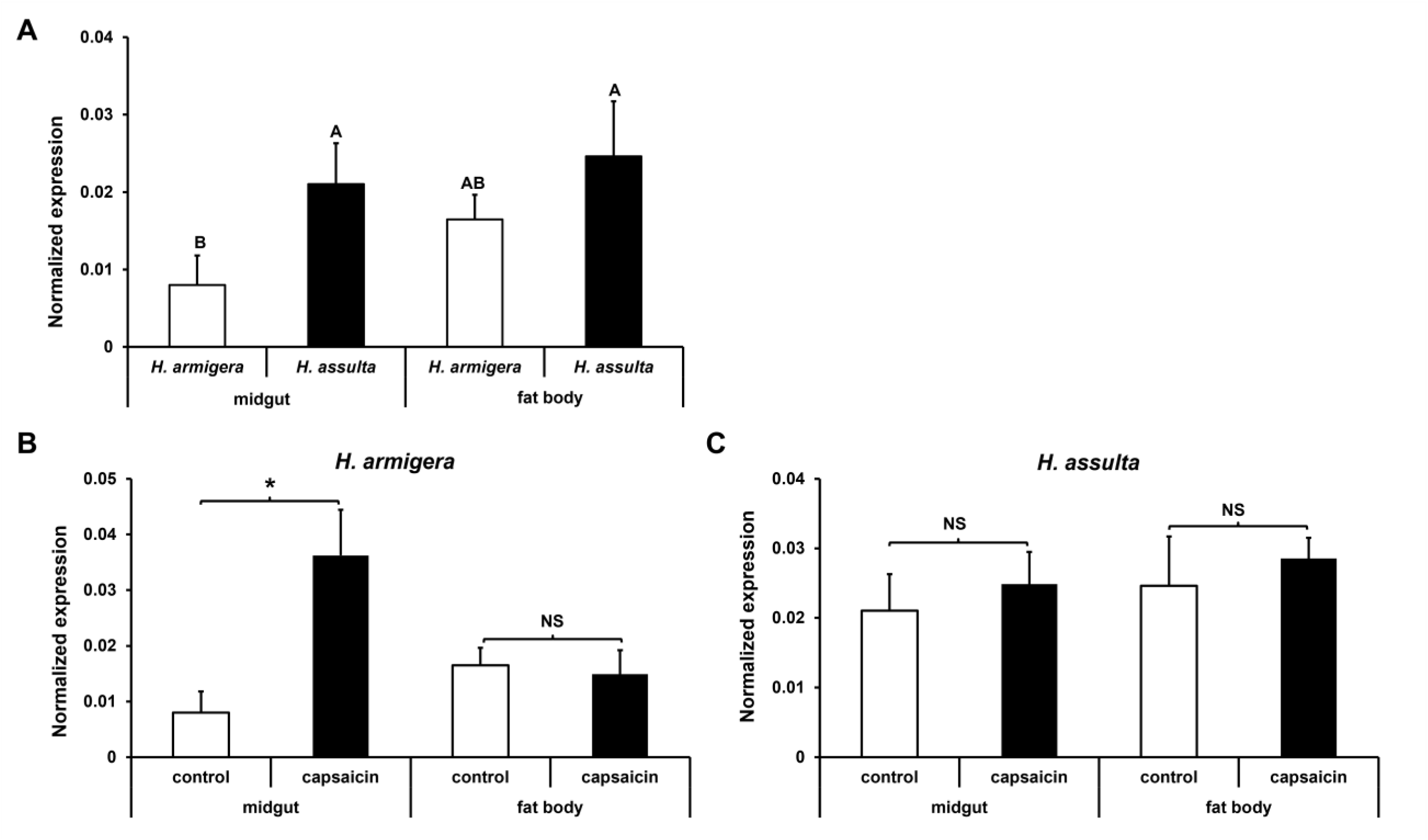
Constitutive and capsaicin-induced *CYP6AN1* expression in the midgut and fat body of *H. armigera* and *H. assulta*. A: Data are presented as mean and standard deviation based on three biological replicates, each consisting of 10 midguts or fat bodies from larvae fed a normal diet. Different letters above the bars indicate significant differences at *p* < 0.05 (one-way ANOVA followed by Tukey′s HSD tests). B and C: Data are presented as mean and standard deviation based on three biological replicates, each consisting of 10 midguts or fat bodies from larvae fed control diet (methanol) or capsaicin-treated diet for 48 h. Significant differences between methanol- and capsaicin-treated larvae in the midgut or fat body are indicated by asterisks (*p* < 0.05, Student′s *t*-test). NS indicates no significant difference.

### 3.4 Recombinant enzyme validation and activities toward model substrates

NADPH–cytochrome *c* reduction assays detected significant reductase activity in all four crude enzyme preparations from *E. coli* cells co-transformed with the respective pCWori-CYP6AN1 construct and pACYC184-HarmCPR. In contrast, no reductase activity was detected in the empty-vector control containing pACYC184 and pCWori (Fig. S2A). Reduced CO-difference spectroscopy revealed a characteristic absorbance peak at 450 nm only in preparations expressing ompA-HarmCYP6AN1 or 17α-HassCYP6AN1, confirming the presence of correctly folded, functional P450 protein (Fig. S2C and F). Accordingly, these two preparations were used in subsequent enzyme assays.

Activities toward the model P450 substrates 7-ethoxycoumarin and 7-methoxyresorufin were evaluated using ECOD and MROD assays, respectively. The HarmCYP6AN1 and HassCYP6AN1 preparations exhibited ECOD activities of 94.78 ± 4.33 and 67.42 ± 1.04 pmol/min/pmol P450, respectively, and MROD activities of 93.35 ± 0.81 and 79.91 ± 2.70 pmol/min/pmol P450, respectively. HarmCYP6AN1 showed significantly higher activity than HassCYP6AN1 toward both model substrates (Table 1). Only background-level ECOD and MROD activities were detected in reactions containing the empty-vector preparation or heat-inactivated enzyme, or in reactions lacking NADPH (data not shown). Together with the reduced CO-difference spectroscopy, these results confirmed that both recombinant CYP6AN1 proteins were catalytically functional in the in vitro system.

**Table 1.**
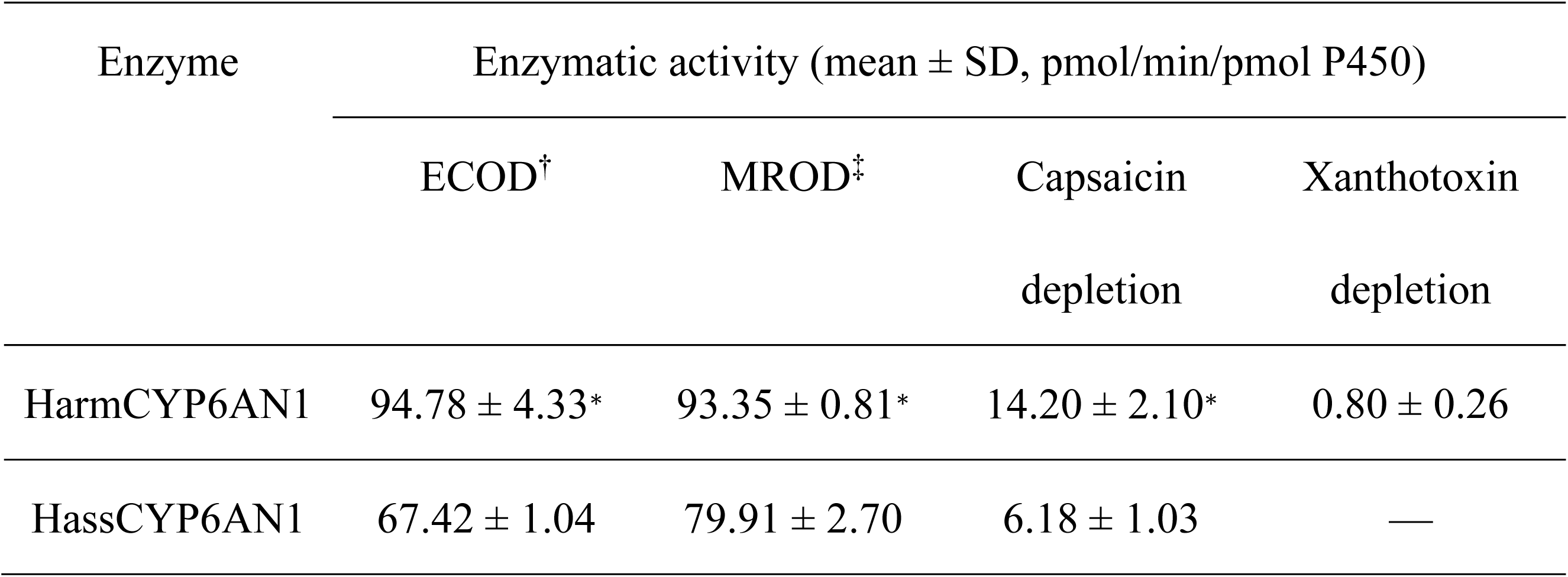
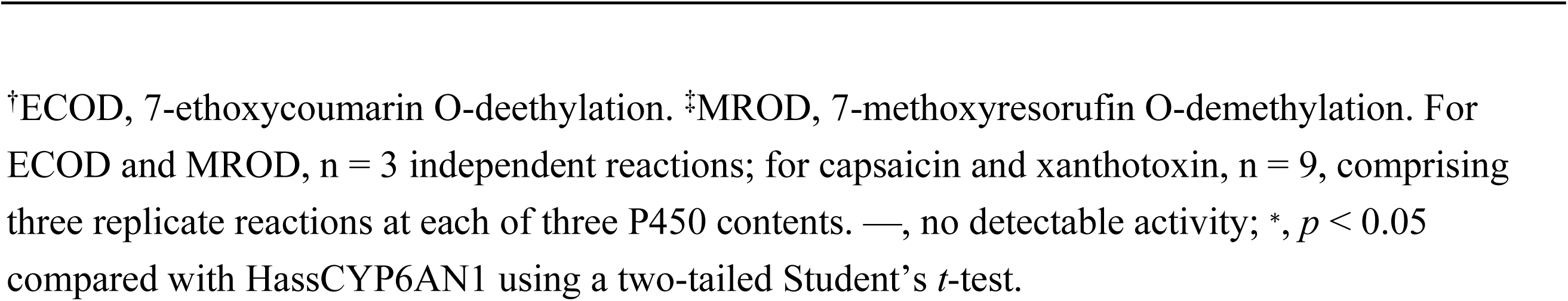
Enzymatic activities of HarmCYP6AN1 and HassCYP6AN1 toward two model substrates and two plant phytochemicals.

| Enzyme | Enzymatic activity (mean ± SD, pmol/min/pmol P450) | | | |
| --- | --- | --- | --- | --- |
|  | ECOD<sup>†</sup> | MROD<sup>‡</sup> | Capsaicin depletion | Xanthotoxin depletion |
| HarmCYP6AN1 | $94.78 \pm 4.33^*$ | $93.35 \pm 0.81^*$ | $14.20 \pm 2.10^*$ | $0.80 \pm 0.26$ |
| HassCYP6AN1 | $67.42 \pm 1.04$ | $79.91 \pm 2.70$ | $6.18 \pm 1.03$ | — |

<sup>†</sup>ECOD, 7-ethoxycoumarin O-deethylation. <sup>‡</sup>MROD, 7-methoxyresorufin O-demethylation. For ECOD and MROD, n = 3 independent reactions; for capsaicin and xanthotoxin, n = 9, comprising three replicate reactions at each of three P450 contents. —, no detectable activity; \*, $p < 0.05$ compared with HassCYP6AN1 using a two-tailed Student's *t*-test.

### 3.5 In vitro metabolism of capsaicin and xanthotoxin

Capsaicin and xanthotoxin metabolism was assessed in NADPH-dependent in vitro reactions using membrane fractions containing recombinant HarmCYP6AN1/HarmCPR or HassCYP6AN1/HarmCPR, and substrate depletion was quantified by HPLC across increasing P450 contents. The amounts of capsaicin remaining decreased with increasing functional P450 content for both HarmCYP6AN1 and HassCYP6AN1 in reactions without PBO, whereas they remained largely unchanged in the corresponding PBO-treated controls (Fig. 5A), consistent with the well-established inhibitory effect of PBO on P450 activity (Casida, 1970). The resulting capsaicin depletion increased with increasing P450 content for both CYP6AN1 enzymes (Fig. 5C). HarmCYP6AN1 depleted capsaicin at an average rate of 14.20 ± 2.10 pmol/min/pmol P450, which was significantly higher than the rate of 6.18 ± 1.03 pmol/min/pmol P450 observed for HassCYP6AN1 (Table 1). Thus, both enzymes metabolized capsaicin, but HarmCYP6AN1 exhibited an approximately 2.3-fold higher depletion rate under the conditions tested. For xanthotoxin, the amount of substrate remaining decreased with increasing HarmCYP6AN1 content in reactions without PBO but remained relatively constant in the PBO-treated controls. In contrast, no consistent P450-content-dependent decrease was observed for HassCYP6AN1 (Fig. 5B). Accordingly, xanthotoxin depletion increased with increasing P450 content only for HarmCYP6AN1, whereas no P450-content-dependent depletion was detected for HassCYP6AN1 (Fig. 5D). HarmCYP6AN1 depleted xanthotoxin at an average rate of 0.80 ± 0.26 pmol/min/pmol P450 (Table 1), indicating that detectable xanthotoxin metabolism was restricted to the *H. armigera* ortholog under the assay conditions.

**Fig. 5.**
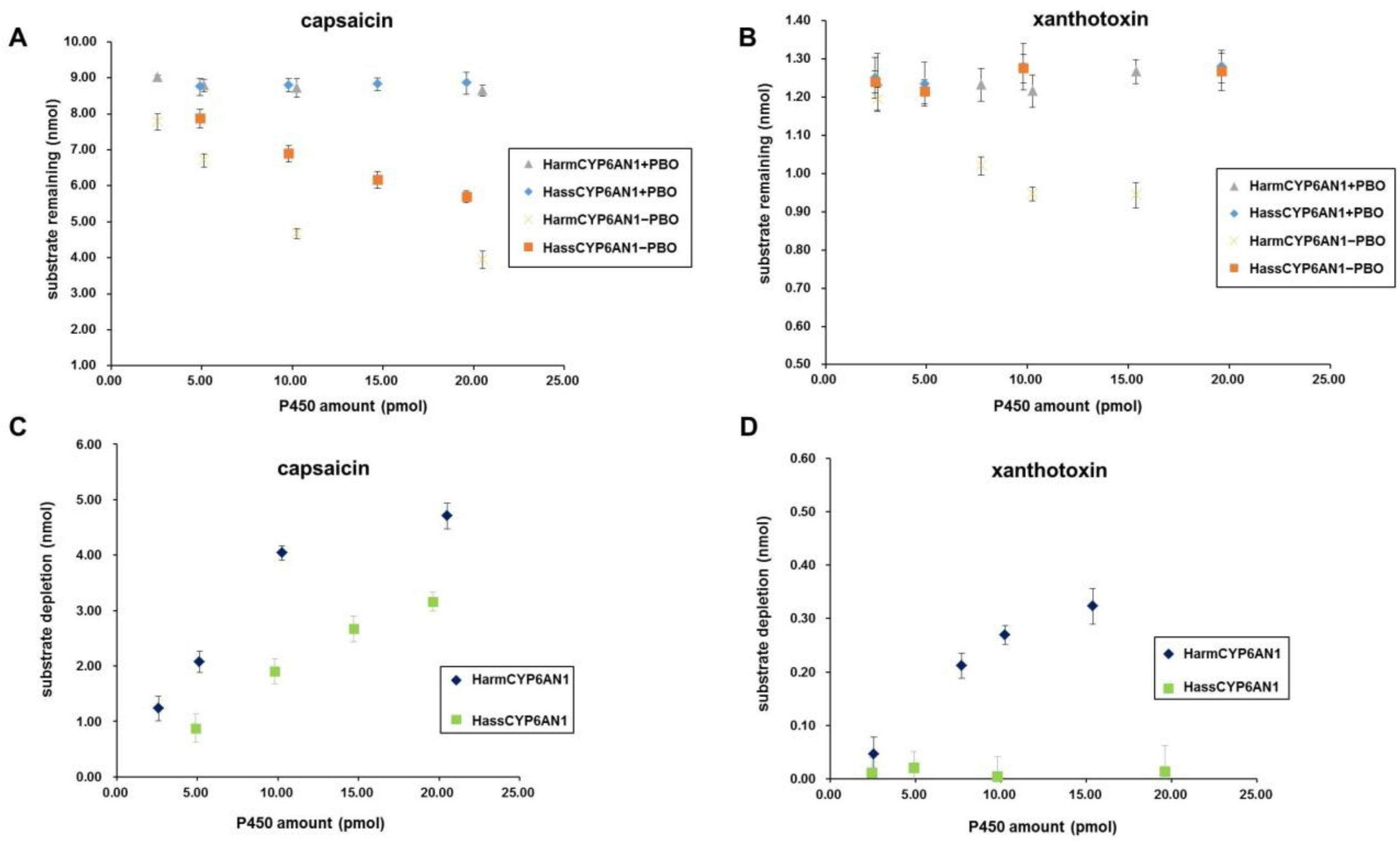
P450-content-dependent metabolism of capsaicin and xanthotoxin by recombinant HarmCYP6AN1 and HassCYP6AN1. A and B: Amounts of capsaicin (A) and xanthotoxin (B) remaining after incubation with increasing amounts of functional CYP6AN1 co-expressed with HarmCPR in *Escherichia coli* membrane preparations. Reactions were conducted in the presence (+PBO) or absence (−PBO) of piperonyl butoxide, with PBO-treated reactions serving as P450-inhibited controls. C and D: P450-dependent depletion of capsaicin (C) and xanthotoxin (D), calculated by subtracting the amount of substrate remaining in the −PBO reaction from that remaining in the corresponding +PBO control. Points and error bars represent the mean ± SD of three independent reactions at each P450 amount.

## 4. Discussion

Comparisons of detoxification genes between closely related generalist and specialist herbivores often emphasize gene-family expansion or gene loss, but evolutionary differentiation can also occur within retained one-to-one orthologs through conservation of a core metabolic function alongside changes in gene regulation and catalytic properties. The *CYP6AN1* orthologs provide an example of this pattern in the sibling species *H. armigera* and *H. assulta*. Both orthologs retain the capacity to metabolize capsaicin, but they differ substantially in transcript organization and expression profiles, as well as in the magnitude and breadth of their detectable catalytic activities. This contrast suggests that adaptation to different host-use strategies may involve changes in how an ancestral detoxification function is deployed rather than its complete replacement.

The assignment of HarmCYP6AN1 and HassCYP6AN1 as orthologs was supported by their high sequence similarity (Fig. 2) and placement within the same well-supported phylogenetic lineage (Fig. 3). Establishing this relationship is important because it indicates that the observed differences in transcription, expression, and catalytic properties arose within a conserved CYP6AN1 lineage rather than between more distantly related CYP6 proteins. Within this lineage, the two species differed markedly in *CYP6AN1* transcript organization. Manual examination of the 5′ flanking region of *HassCYP6AN1* identified three putative TATA-box-associated promoter regions, P1, P2, and P3 (Fig. 1 and S1). The RACE results suggest that alternative transcription initiation at these regions, together with alternative splicing within the 5′ UTR, generates transcripts with distinct 5′ UTR structures but an identical *HassCYP6AN1* coding sequence. In contrast, the *HarmCYP6AN1* upstream region lacks the TATA-box elements corresponding to P2 and P3 (Fig. S1), consistent with the detection of only a single full-length transcript, although the causal contribution of these elements remains to be tested. Because the *HassCYP6AN1* isoforms encode the same protein, any functional differences among them would be expected to involve transcript abundance, stability, localization, or translational efficiency rather than the intrinsic catalytic properties of the encoded protein. Although comparable information from insects remains limited, alternative 5′ UTRs in mammals generally evolve more rapidly and experience weaker purifying selection than coding regions (Resch *et al*., 2009) and can influence transcript stability and translational efficiency (Bicknell *et al*., 2012; Kramer *et al*., 2013). Both *CYP6AN1* genes also contain introns within their 5′ UTRs, providing an additional potential layer of regulation because such introns can affect gene expression, transcript localization, and translational control (Cenik *et al*., 2010; Bicknell *et al*., 2012). Thus, the combination of multiple putative promoters, alternative 5′ UTRs, and 5′-UTR introns may provide *HassCYP6AN1* with greater regulatory flexibility.

More broadly, the pronounced divergence in 5′ regulatory architecture, despite strong conservation of the coding region and capsaicin-metabolizing function, suggests that evolutionary differentiation of *CYP6AN1* has involved substantial regulatory modification without loss of its shared metabolic role. Such regulatory changes may allow this conserved detoxification function to be deployed differently in the two species. However, isoform-specific expression analyses and functional promoter assays will be required to establish the biological significance of these differences.

The differences in regulatory architecture were accompanied by contrasting expression profiles of the two *CYP6AN1* orthologs. *HassCYP6AN1* maintained higher constitutive expression, particularly in the larval midgut, whereas *HarmCYP6AN1* was strongly induced by dietary capsaicin (Fig. 4). Generalist herbivores encountering chemically variable diets have been proposed to rely more heavily on inducible detoxification responses, whereas specialists repeatedly exposed to a narrower range of host-associated compounds may maintain higher constitutive expression of relevant detoxification enzymes (Li *et al*., 2007). Viewed within this framework, the relatively high constitutive expression of *HassCYP6AN1* may be associated with the more restricted host use of specialist *H. assulta*, whereas the inducibility of *HarmCYP6AN1* may provide a flexible response to the chemically diverse diet of generalist *H. armigera*. Whether the more complex promoter and transcript architecture of *HassCYP6AN1* contributes directly to its higher constitutive expression remains unresolved. Moreover, because the expression assays quantified total *HassCYP6AN1* transcripts, the constitutive abundance and capsaicin responsiveness of individual isoforms require further investigation.

Recombinant enzyme assays demonstrated that both CYP6AN1 orthologs metabolized capsaicin. Capsaicin depletion increased with increasing functional P450 content in both enzyme preparations (Fig. 5C), whereas no comparable P450-content-dependent depletion was observed in the PBO-treated controls (Fig. 5A), supporting the conclusion that capsaicin metabolism depended on an active CYP6AN1/CPR system. Under the conditions tested, HarmCYP6AN1 exhibited a capsaicin-depletion activity of 14.20 ± 2.10 pmol/min/pmol P450, approximately 2.3-fold higher than that of HassCYP6AN1 (6.18 ± 1.03 pmol/min/pmol P450). To place these activities in context, depletion rates were normalized to functional P450 content following Tian *et al*. (2019), who characterized several capsaicin-metabolizing P450s from *H. armigera* using a similar *E. coli*-based expression system. The value for HarmCYP6AN1 was numerically higher than those reported for CYP9A12 (11.9 ± 1.5 min^-1^, equivalent to pmol/min/pmol P450) but lower than those for CYP9A17 (14.8 ± 0.2 min^-1^), CYP9A14 (16.3 ± 1.4 min^-1^), and CYP6B6 (21.9 ± 0.1 min^-1^); HassCYP6AN1 showed lower activity than all four enzymes. Thus, HarmCYP6AN1 falls within the activity range of previously characterized capsaicin-metabolizing P450s, whereas HassCYP6AN1 displays comparatively lower activity under the assay conditions used. The higher capsaicin-depletion activity of HarmCYP6AN1 was accompanied by higher ECOD and MROD activities, consistent with greater catalytic activity toward multiple tested substrates rather than a difference restricted to capsaicin. Nevertheless, CYP6AN1 alone is unlikely to determine species-level capsaicin tolerance because capsaicinoid metabolism in *Helicoverpa* involves multiple P450s and UDP-glycosyltransferases (Ahn *et al*., 2011; Tian *et al*., 2019; Xiong *et al*., 2025). Moreover, normalized depletion activities are assay-dependent and should not be interpreted as intrinsic catalytic efficiencies equivalent to k_cat_/K_m_. Determination of K_m_, V_max_, product identities, and reaction stoichiometry will be required to define the kinetic and biochemical basis of the observed differences.

The xanthotoxin assay revealed a further substrate-specific difference between the two orthologs. HarmCYP6AN1 showed P450-content-dependent xanthotoxin depletion, whereas HassCYP6AN1 showed no consistent increase in depletion with P450 content (Fig. 5D), indicating that xanthotoxin metabolism was detectable only for HarmCYP6AN1 under the conditions tested. Xanthotoxin is a plant furanocoumarin encountered by polyphagous herbivores and is metabolized by several *H. armigera* P450s, including CYP6AE19 and members of the CYP9A subfamily (Wang *et al*., 2018; Tian *et al*., 2024). The ability of HarmCYP6AN1 to metabolize both capsaicin and xanthotoxin is consistent with its participation in a detoxification network that processes chemically distinct plant compounds. However, the absence of detectable xanthotoxin metabolism by HassCYP6AN1 should not be interpreted as evidence of reduced overall catalytic competence, because the enzyme retained activity toward both model substrates and capsaicin. Instead, the results indicate a narrower detectable substrate range under the experimental conditions used, and HassCYP6AN1 may metabolize other host-associated compounds not examined in this study.

The high amino acid sequence similarity between HarmCYP6AN1 and HassCYP6AN1 provides an important context for interpreting their catalytic differences. The two proteins share 97.5% identity and differ at 13 of 518 positions, including two substitutions within predicted substrate recognition sites (SRSs) (Fig. 2). Because substitutions within SRSs can alter P450 substrate recognition and metabolism, these residues are plausible candidates for contributing to the higher capsaicin-depletion activity and detectable xanthotoxin metabolism of HarmCYP6AN1. Consistent with this possibility, Shi et al. (2022) showed that a single amino acid substitution within SRS4 of *H. armigera* CYP6AE20 conferred activity toward xanthotoxin. However, sequence comparison alone does not establish a mechanistic basis for the observed differences, and substitutions outside the SRSs may also affect protein conformation, substrate access, or catalytic activity. Reciprocal site-directed mutagenesis followed by normalized recombinant enzyme assays will therefore be required to determine which substitutions contribute to the catalytic differences between the two orthologs. Thus, HarmCYP6AN1 and HassCYP6AN1 retain a conserved capacity for capsaicin metabolism while differing in activity magnitude and detectable substrate range.

Taken together, these findings establish CYP6AN1 as a component of the capsaicinoid-metabolizing repertoire in both *H. armigera* and *H. assulta*, while showing that the two species differ in how this ortholog is regulated and functions. *H. assulta* combines higher constitutive expression with greater 5′-UTR transcript diversity, whereas *H. armigera* combines capsaicin-inducible expression with higher capsaicin-depletion activity and detectable activity toward xanthotoxin. These differences illustrate how regulatory and catalytic variation can arise within an orthologous detoxification lineage while preserving metabolism of a shared host-plant compound. CYP6AN1 should therefore be regarded as one component of the multienzyme system involved in capsaicinoid metabolism rather than as the sole determinant of capsaicin metabolism or tolerance. The contrasting regulation and detectable substrate ranges of these orthologs provide a useful system for investigating how conserved detoxification functions vary between closely related herbivores with contrasting diet breadths.

## Acknowledgments

This work was supported by the National Natural Science Foundation of China (NSFC)–Henan Joint Major Grant (No. U2004206), an NSFC grant (No. 31672350), and a Swanlund Chair Endowment to M. R. Berenbaum. We also thank Lihong Yang for insect rearing and Xiongya Wang for assistance with tissue dissection during the experiments.

## Data availability statement

The data supporting the findings of this study are available in the Illinois Data Bank (https://databank.illinois.edu/datasets/IDB-8268315?code=lXgHdcuYzayzYst-a98Wkp-9zoBlF-L6RXwayftN764).

---

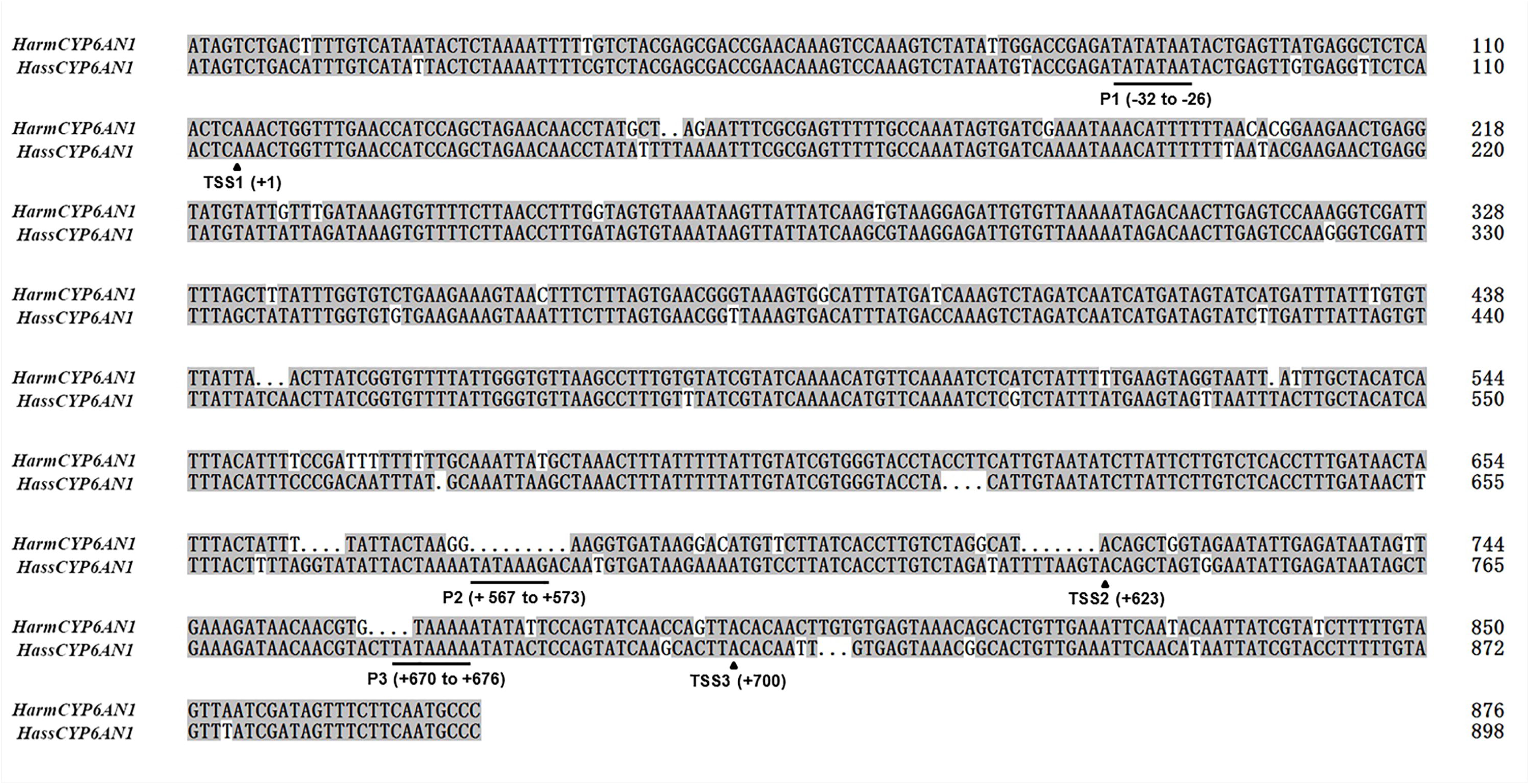

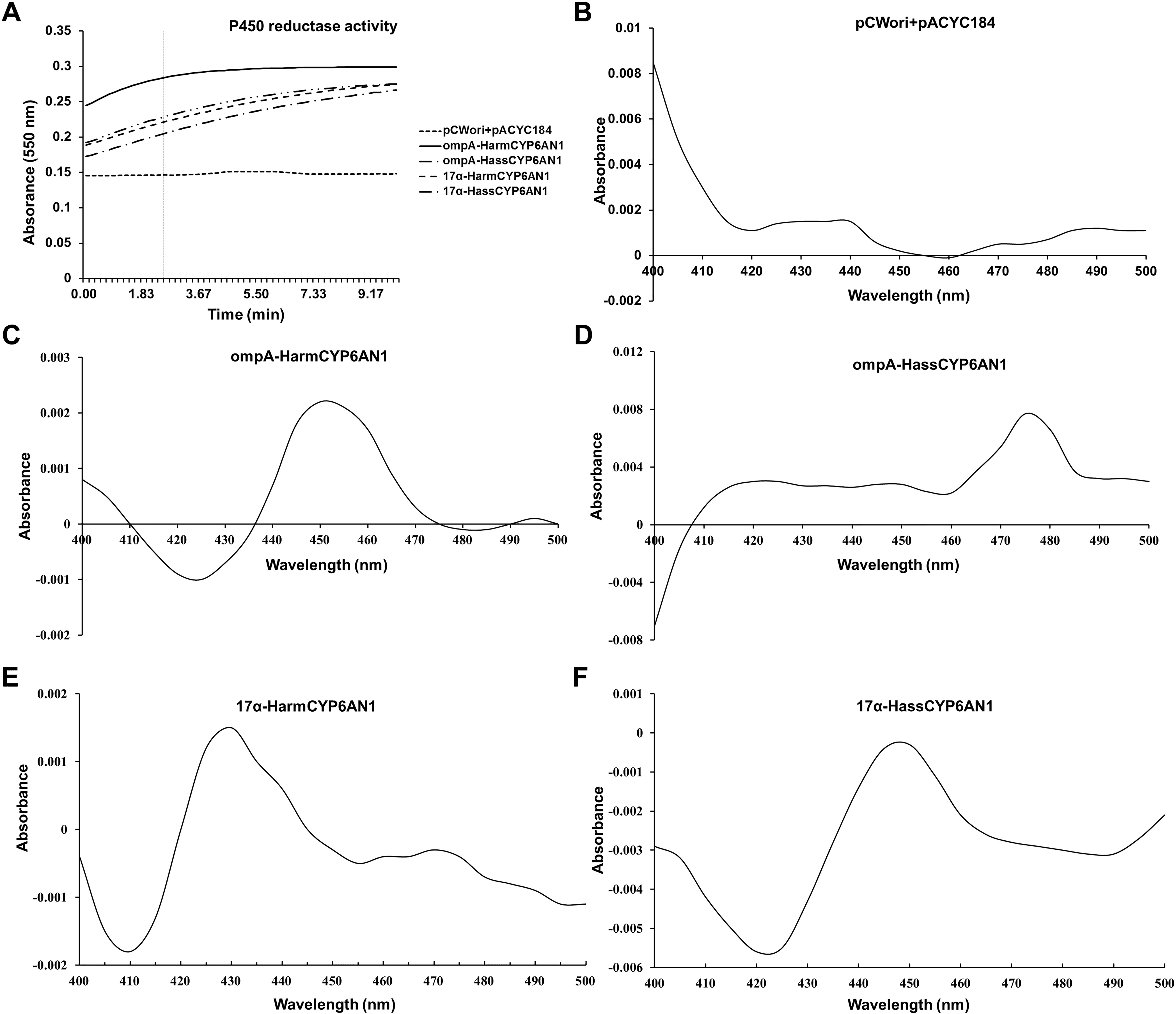

