## Supplementary figures and tables for "Molecular and functional characterization of CYP6AN1 orthologs involved in capsaicin metabolism in two *Helicoverpa* species"

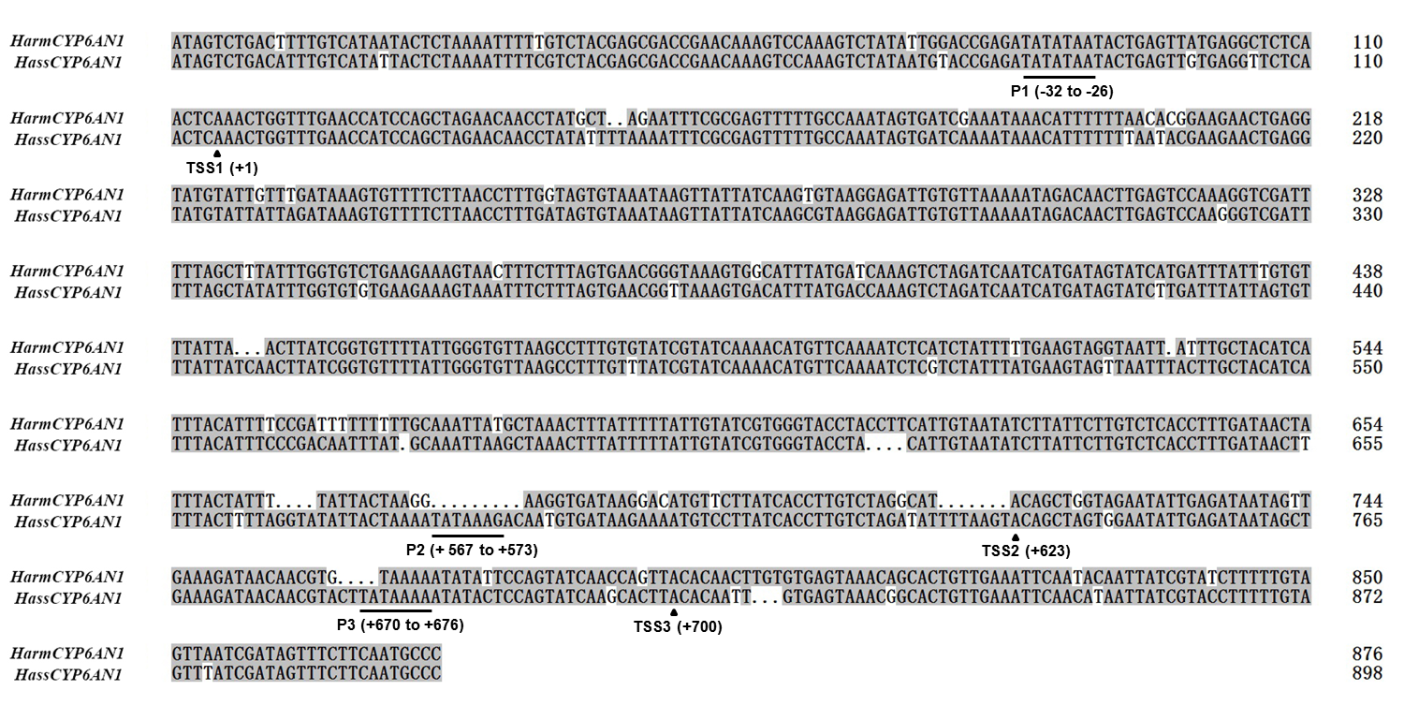

**Fig. S1.** **Sequence comparison of the 5′ flanking regions of *HarmCYP6AN1* and *HassCYP6AN1***. The sequences represent positions −114 to +762 of the 5′ flanking region of *HarmCYP6AN1* and −114 to +784 of the 5′ flanking region of *HassCYP6AN1*. The TATA-box elements with relative positions in the *HassCYP6AN1* promoters are underlined. The corresponding transcription start sites (TSSs) and their relative positions in each promoter are indicated by arrows.

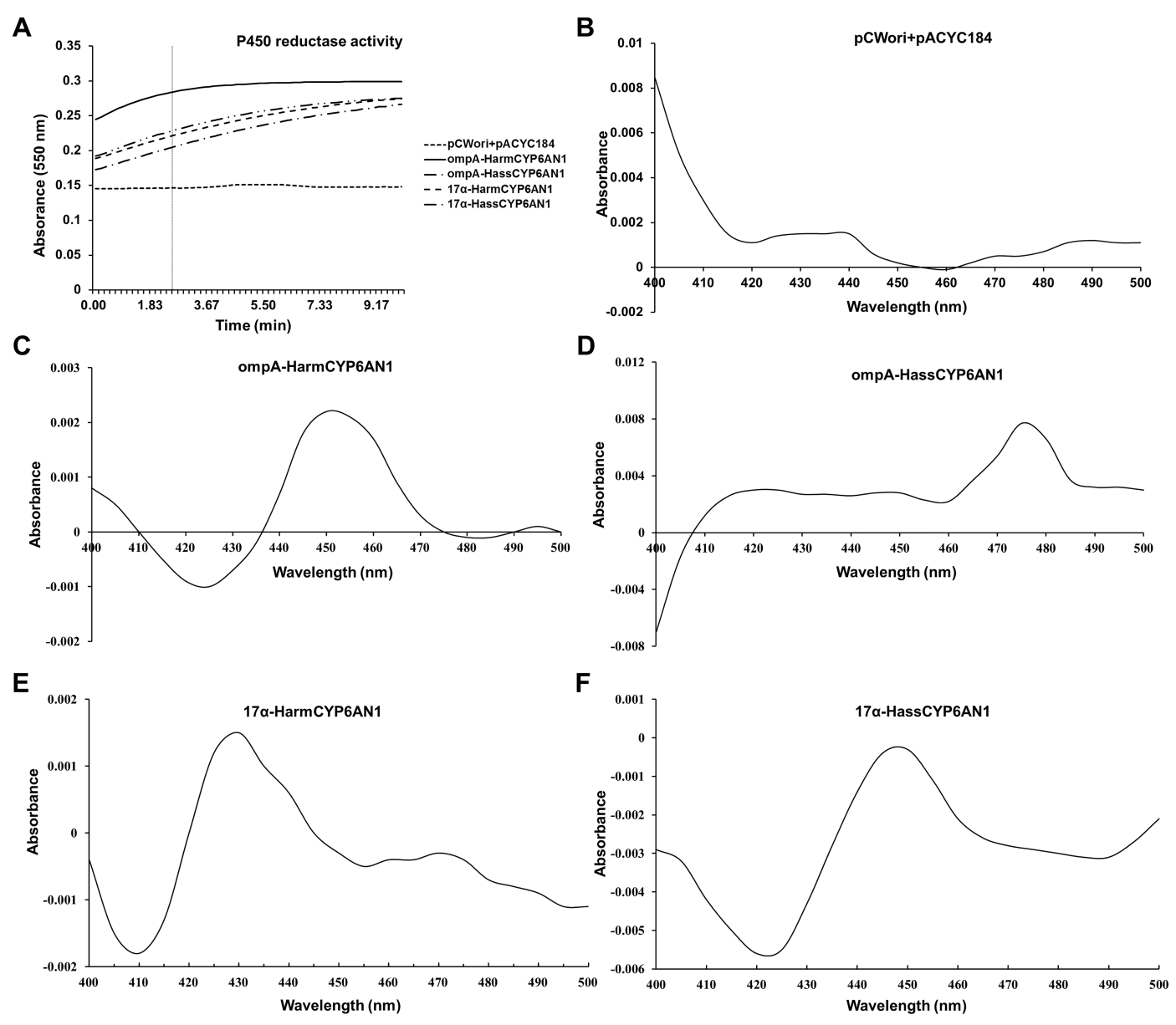

**Fig. S2. Enzymatic activity of *E. coli*–expressed HarmCPR and reduced CO-difference spectra of the noninsertion control and recombinant CYP6AN1 variants**. A: Data represent mean values from three independent measurements at each time point. The vertical black dashed line indicates the specific time point (2.57 min) used for calculating the enzymatic activities. B-F: ompA and 17α correspond to proteins expressed using the ompA+2 and 17α strategies, respectively. pCWori + pACYC184 indicates expression via co-transformation of two noninsertion control vectors into *E. coli*. Each panel presents a representative spectrum from multiple scans.

**Table S1. PCR primers used in this study.**

| Primer name | Sequence (5′-3′) | Purpose |
| --- | --- | --- |
| UPM | CTAATACGACTCACTATAGGGCAAGCAGTGGTATCAACGCAGAGT | RACE characterization of *CYP6AN1* |
| UPS | CTAATACGACTCACTATAGGGC |  |
| R-GSP | AGGGTTTCTGCCGATTCCTCTTGGGT |  |
| R-NGSP | CACAACTTTCTCATTGGGATACTGGCGG |  |
| F-GSP | CGAGAAATATTTTGAACATCCGGAGAAGTTC |  |
| F-NGSP | GTGAAGGACCTCGGCAGTGC |  |
| 3′RP | GCTGTCAACGATACGCTACGTAACG |  |
| 3′NRP | CGCTACGTAACGGCATGACAGTG |  |
| M-F | TCTCAAACTGGTTTGAACCATC | Cloning full-length cDNA and gDNA of *CYP6AN1*s |
| M-R | TTAAATTGTATCTTATTTATTTTAAGTGCTTAAAAT |  |
| S-F1 | AAACTGGTTTGAACCATCCAG |  |
| S-F2 | AGTACATGGACACAATTGTGAG |  |
| S-F3 | TACATGGACACAATTGTGAGTAAAC |  |
| S-R | TTGCGTTTTATAAAAAATTATCTTATATATTTTTAGTGC |  |
| qPCR-F | GGTAGTTGGGTGCGCTGAGA | RT-qPCR assay of *CYP6AN1* expression |
| qPCR-R | ATCTGTATGCCGAAGCCGCA |  |
| EF-1α-F | GAAGTCAAGTCCGTGGAGATG |  |
| EF-1α-R | GACCTGTGCTGTGAAGTCG |  |
| RPL32-F | CATCAATCGGATCGCTATG |  |
| RPL32-R | CCATTGGGTAGCATGTGAC |  |
| ompA-F (Nde I) | GGAATTCCATATGAAAAAGACAGCTATCGCGATTGCAGTGGCACTGGCTGGTTTCGCTA | Constructing pCWori-*CYP6AN1* plasmid |
| ompA-F (BamH I) | CGCGGATCCATGAAAAAGACAGCTATCGCGATTGCAGTGGCACTGGCTGGTTTCGCTAC |  |
| M-L | AGTAGCAGCAGGAATATCATCGGAGCGGCCTGCGCTACGGTAGCGAAACCAGCCAGTGC |  |
| M-17α-F (Nde I) | GGAATTCCATATGGCTCTGTTATTAGCAGTTTTTCTCGTGCTCTTATACAAATACAC |  |
| S-L | AGTAGCAGAAGGAATATCATCGGAGCGGCCTGCGCTACGGTAGCGAAACCAGCCAGTGC |  |
| S-17α-F (BamH I) | CGCGGATCCATGGCTCTGTTATTAGCAGTTTTTCTCGTGCTCTTATACTACTAC |  |
| M-R (Hind III) | CCCAAGCTTTTATGCAGCAACATTATTTTCCT |  |
| S-R (Hind III) | CCCAAGCTTTTATGCAGCCACATTATTTTC |  |
| pelB-F (Nde I) | GGAGATATACATATGAAATACCTGCTGCCGACCGCTGCTGCTGGTCTGCTGCTCCTC | Constructing pACYC184-HarmCPR plasmid |
| CPR-L | TCCTGTGCGCTGTCTGACATGGCCATCGCCGGCTGGGCAGCGAGGAGCAGCAGACCAGC |  |
| CPR-R (Hind III) | CCCAAGCTTTTAACTCCATACATCTGCTGAATATTTCTTCATAGACTC |  |
| CPR-homoF | CGATGCTTAGGAGGTCATATGAAATACCTGCTGCCGAC |  |
| CPR-homoR | CAGCTTATCATCGATAAGCTTTTAACTCCATACATCTGCTG |  |
| pCWori-homoF | CCACACCCGTCCTGTGGATCCATTCGATGGTGTCCTGG |  |
| pCWori-homoR | TGCGTCCGGCGTAGAGGATCCGCGTACTATGGTTGCTTTGAC |  |
