## Supplementary file 1 for "Molecular and functional characterization of CYP6AN1 orthologs involved in capsaicin metabolism in two *Helicoverpa* species"

> *H. armigera* CYP6AN1 (GenBank: XIF13453.1)

MIFLLLLPIALVLLYKYTTRNHDYWEKRNIKHEKPIPVFGTLYDNVFAKKSIIEVAADLYRQYPNEKVVGMFRGNTPELVIRDLEIVRKILNVDFAYFYPRGIGRNPTYEPLFMNLFHAEGDSWKLLRQRLTSTFTTAKLKKMFPLVVGCAEKLTHVGEDIVNKGGDCDVRELMARFTTEFIGACGFGIQMDTINDEHSLFRDLGRKMFNRSTKIHFLVALYDLFPEFGTFFQKTLKETFLLDSMTEIVQNIRKQRNFKPIGRNDFIDLLLELEEKGKITGESVEKRTADGKPVQVEMEMDLTCMVAQVFVFFAAGFETSSSATSFMLHQLAFHPEEQKKIQEDIDRVLAKYNNQLCYDSISEMKALSNGFKEAMRMFPSLGTLHRVCAKKYKIEELGITIDPDVKIIIPLQGIQNDEKYFEHPEKFMPDRFNDTSEERNKFAYMPFGEGPRQCIGARLGEMQSLAGLAAVLHKFSVEPTTTTKRYPDINHGSNIVQSIEGGLPLKLKLRSKENNVAA

> *H. armigera* CYP6AN3 (GenBank: WRX06083.1)

MLLLLIPIVLVLLYYYTTRNHDYWEKRNIKHEKPIPVFGTLYNNVFAKKSITEISVELYRQYPNEKVVGIYRGTTPELIIRDLDIARKILNVDFAYFYPRGLGRNPKYEPAFLNIFHVDGDTWKLLRQRLTSAFTTAKLKSMFPLVVACAEKLQNVGEDIVNKGGDCDVRELMARFTTEFIGACGFGIQMDTINNEHSLFRDLGRKMFNRSWRNFVLVPLYDLFPDFRTLFQKILHEPFLLDSMAEIVQNIRKQRNNKPIGRNDFIDLLLEVEAKGKMTGESVEKRTADGKPVQVEMDMDLTCMVAQVFVFFAAGFETSSSATSFLLHQLAFHPEEQKKIQEDIDRVLAKYNNQLCYDSISEMKALSNGFKEAMRMFPSLGTLHRVCAKKYKIEELGITIDPDVKIIIPVEGIQNDEKYFENPTQFKPDRFNDPSEERHKFAYMPFGEGPRQCIGARLGEMQSLAGLAAVLQKFSVEPAATTKRYPEVNHGSNTVQSIKGGLPLKLTLRNK

> *H. armigera* CYP6AE14 (GenBank: WRX06019.1)

MITSLLLTAVFVIIFTIYLVSKKKFQYWEKRKVPHLPPVPLLGNFGNFILQRQFLGYTLQQICGKFPNVPYVGAYFGTEPALIVQDPEHIKLVMTKDFYFFSSREISEYADRERFTQNLFSTSGNKWKVLRQNLTPVFTSAKMKNMFHLIEKCSHVFEDFLDKEAKSNEVEMRALVARYTMDCIGTCAFGVETKTMNVTENNPFTAVGNSIFMLSRVQGFKFVLRGIYPSLFYLLGFRTLPPEVNAFFSNLMTGVFKGRNYAPTSRNDFVDFVLKWKQNKTMTGDSLTNMKYDSQKKVTLEVDDDLLVAQCFIFFAAGYETSATTLSFTLYELAKHPEAQKRAIAEVDDYLRRHNNELKYECLSEMPFVEACFDETLRKYPVLSLLTREVVEDYTFPSGLKVEKGLRIFLPLYHLHHNPEFFPDPEEYRPERFLPENKDKIKPYTYMPFGDGPRLCIGMRFAKMQMTAGIITLLKKYRLELAPGMPQNIEFEPNSFVSQVAGGINLKMIKRESWEGRLLKNLEKAY

> *H. armigera* CYP6AE16 (GenBank: AID54891.1)

MITLLLLFAVFSIISAIYLLSKKKFQYWEKKKVPHPPPVPLLGNFGNYILQREFLGDIVKEICDKFPNAPYIGAYYATEPALIVKDPELIKLVITKDFYFFSNREVAEFSNREKLTQNLFATSGNQWKVLRQNLTPVFTSAKMKNMFHLIEKCSHVFEDFLDKEAKSNEIEMRALIARYTMDCIGTCAFGVDTSTMDKTENNPFTAVGNSIFILDWFQSFKFTLRSIYPSIFYALRYTTLPPMVEAFFSNLMTGVFKGRNYTPTSRNDFVDFVLKWKHDKTITGDSLENLKSNSQKKVSLEVDDDLLVAQCFIFFAAGYETSATTLSFTLYELAKNPEAQKRAIAEVDDYLRRHNNELKYECLSEMPFVEACFDETLRKYPVLSVLAREVVEDYTFPSGLKVEKGLRILLPLYHLHHNPEFFPDPEEYRPERFLPENKHNIKPYTYMPFGDGPRLCIGMRFAKMQMTAGIITLLKKYRLELAPGMKTKVEFQPKSMVTQVIGGIKLKMIEREGWEERLLKKL

> *H. assulta* CYP6AN1 (GenBank: XIF13454.1)

MIFLLLLPIALVLLYYYTTRNHDYWEKRNVKHQKPIPVFGTLYDNVFAKKSIIEMAADLYRQYPNEKVVGMFRGNTPELVIRDLEIVRKILNVDFAYFYPRGIGRNPTYEPLFMNLFHAEGDSWKLLRQRLTSTFTTAKLKKMFPLVVGCAEKLTHVGEDIVNKGGDCDVRELMARFTTEFIGACGFGIQMDTINDEHSLFRHLGRKMFNRSTKIHILVALYDLFPELGTFFQKTLRETFLLDSMTEIVQNIRKQRNFKPIGRNDFIDLLLELEGKGKITGESVEKRTADGKPVQVEMEMDLTCMVAQVFVFFAAGFETSSSATSFMLHQLAFHPEEQKKIQEDIDRVLAKYNNQLCYDSISEMKALSNGFKEAMRMFPSLGTLHRVCAKKYKIEELGITIDPDVKIIIPLQGIQNDGKYFEHPEKFLPDRFNDASEERNKFAYMPFGEGPRQCIGARLGEMQSLAGLAAVLHKFSVEPTTTTKRYPDINHASNIVQSIEGGLPLKLKLRSKENNVAA

> *H. assulta* Chr22 partial (genome-derived, GCA_029618815.1, chromosome 22)

YYYTTRNHDYWEKRNVKHEKPIPVFGTLYNNVFAKKSITEISVELYRQYPNEKVVGIYRGTTPELIIRDLDIARKILNVDFAYFYPRGLGRNPKYEPAFLNIFHVDGDTWKLLRQRLTSAFTTAKLKSMFPLVVACAEKLQNVGEDIVSRGGDCDVRELMARFTTEFIGACGFGIQMDTINNEHSLFRDLGRKMFNRSWRNFVLVPLYDLFPDFRTLFQKILHEPFLLDSMAEIVQNIRKQRNFKPIGRNDFIDLLLEVEAKGKMTGESVEKRTADGKPVQVEMDMDLTCMVAQVFVFFAAGFETSSSATSFLLHQLAFHPEEQKKIQEDIDRVLAKYNNQLCYDSISEMKALSNGFKEAMRMFPSLGTLHRVCAKKYKIEELGITIDPDVKIIIPVEGIQNDEKYFENPTQFKPDRFNDPSEKRHKFAYMPFGEGPRQCIGARLGEMQSLAGLAAVLHKFSVEPAATTKRYPEVNHGSNTVQSIKGGLPLKLTLRNK

> *H. assulta* Chr29 partial (genome-derived, GCA_029618815.1, chromosome 29)

YLYGTRTFDYWKKRGIKHDKPIPIFGTNLRQFLQQASMAMMATEVYNKYPEEKVVGFFRGTAPELVVRDPEIIKRILVTDFQYFHSRGFNPHKTVVEPLLKNIFFADGDLWRLIRQRFTPAFSTGKLKAMFHIITDRAEKLQIITEEVVNKEYYDVRELMARYTTDFIGVCAFGINMDSLNDENSHFRKLGKRIFERRFRDAVAGALKIMFPELCKNMHFLSPELEANMRHLVETVLKNRNYKPSGRNDFIDLMLELRQKGKIIGESIEKKNEDGTSKIVELEMDDLLMTAQAFVFFGAGFETSSTASSYTLHQLAFHPEYQKKVQEEIDTVLAKHNNKLTYDAVKEMHYLEMAFYESMRMYPSVAYLVRMCCVPEYTIPEINLTINQDVKLMIPIQAIHRDEKYFREPHKFDPERFNEGTKEDIKNFVYLPFGEGPRSCVGARLGQMQSMAGLAAVLHKYSVEPAACTVRDPLPEPTGIVSEGFVGGLPLKIRRREK

> *H. zea* P450 6B6-like (1) (GenBank: XP_047019984.1)

MIFLLLLPIALVLLYYYTTRNHDYWEKRNIKHEKPIPVFGTLYDNVFAKKSIIEMAADLYRQYPNEKVVGMFRGNTPELVIRDLEIVRKILNVDFAYFYPRGIGRNPTYEPLFMNLFHAEGDSWKLLRQRLTSTFTTAKLKKMFPLVVGCAEKLTHVGEDIVNKGGDCDVRELMARFTTEFIGACGFGIQMDTINDEHSLFRDLGRKMFNRSTKIHFLVALYDLFPEFGTFFQKTLKETFLLDSMTEIVQNIRKQRNFKPIGRNDFIDLLLELEEKGKITGESVEKRTADGKPMQVEMEMDLTCMVAQVFVFFAAGFETSSSATSFMLHQLAFHPEEQKKIQEDIDRVLAKYNNQLCYDSISEMKALSNGFKEAMRMFPSLGTLHRVCAKKYKIEELGITIDPDVKIIIPLQGIQNDEKYFEHPEKFIPDRFNDTSEERNKFAYMPFGEGPRQCIGARLGEMQSLAGLAAVLHKFSVEPTTTTKRYPEINHGSNIVQSIEGGLPLKLKLRSKENNVAA

> *H. zea* P450 6B6-like (2) (GenBank: XP_047019985.1)

MLLLLIPIVLVLLYYYTTRNHDYWEKRNVKHEKPIPVFGTLYNNVFAKKSITEISVELYRQYPNEKVVGIYRGTTPELIIRDLDIARKILNVDFAYFYPRGLGRNPKYEPAFLNIFHVDGDTWKLLRQRLTSAFTTAKLKSMFPLVVACAEKLQNVGEDIVNKGGDCDVRELMARFTTEFIGACGFGIQMDTINNEHSLFRDLGRKMFNRSWRNFVLVPLYDLFPDFRTLFQKILHEPFLLDSMAEIVQNIRKQRNNKPIGRNDFIDLLLEVEAKGKMTGESVEKRTADGKPVQIEMDMDLTCMVAQVFVFFAAGFETSSSATSFLLHQLAFHPEEQKKIQEDIDRVLAKYNNQLCYDSISEMKALSNGFKEAMRMFPSLGTLHRVCAKKYMIEELGITIDPDVKIIIPVEGIQNDEKYFENPTQFKPDRFNDPSEERHKFAYMPFGEGPRQCIGARLGEMQSLAGLAAVLHKFSVEPAATTKRYPEVNHGSNTVQSIKGGLPLKLTLRKK

> *S. frugiperda* CYP6AN4 (GenBank: AGO62003.1)

MFLLLLPLVLILLYHYVTRNHDYWEKRNVKYEKPVPIFGTIYPNLVAKKSITEIAVELYNKYPNEPAVGIYRGTTPELIVRDLDVVRRILNVDFAYFYPRGIGRNPKEEPVFLNLFHVDGDIWKLLRQRLTPTFTTAKLKNMFPLVVQCAEKLQTVGEDIVNRGGDCDVRELMARFTTEFIGACGFGIQMDSINNEHSLFRALGRNIFHRSWRNYVAVALYDLFPEFRTIIQKSLRETKIMDAIAEIVENIRKQRNYKPIGRNDFIDLLLELESKGKIVGESVEKRDANGKPEQVEMEMDLTCMVAQVFVFFAAGFETSSSATSFMLHQLAYHPEEQRKIQENIDQVLAKYNNKLCYDSISEMTALSNGFKEAMRLFPSLGTLHRVCAQKYTIPEMGITLDPGVKIIVPVQAIQVDGKYFENPTEFLPDRFNDTSADRHKFAYLPFGEGPRQCIGARLGEMQSLAGLAAVLHKFSVEPAANTKRHLEVNHGSNTVQSIRGGLPLKLRLRSKQSHVAA

> *S. frugiperda* P450 6B2 (GenBank: XP_035444666.2)

MFLILVLIGLVALYFYGTRTFNYWKSRGIKHDKPVPIFGTNLKQFMQQASMCMMATEMYRKYPEEKVVGFYRGTDPELVIRDPEIIKRILTTDFQCFHSRGLTYHKTCVEPLLRNIFFADGDLWRLIRQRFTPAFSTGKLKAMFHIITDRAEKLQVITEEVVDREYYDARELMARYTTDFIGVCAFGIDMDSLSDENSHFRKLGKRIFERRFRDAVAGALKIMFPEVFKHMHFLAPELETNMKHLVQTVLKNRNYKPSGRNDFIDLMLELREKGKIIGESIEHKNPDGTPKIVDLEMDDLLMTAQAFVFFGAGFETSSTASSYTLHQLAFHPEYQKKVQEEVDTVLAKHNNKLTYDAIKEMKYLEMAFYESMRMYPSVGYLIRQCCVPKYTFPEIDLTIDEGLKVMIPIQAIHRDEKYFREPNKFDPERFSDGTKEDIKNFVYLPFGEGPRSCVGARLGQMQSMAGLAAVLQKYSVEPAACSRQEPLPDPAGIVSEGFVGGLPLKIRRRVK

> *S. littoralis* CYP6AN4 (GenBank: AFP20585.1)

MSLLLLPLVLVLVYHYVTRNHDYWEKRNVKYEKPVPIFGTLYANLVAKKSITEISVEMYNKYPNEPAVGIYRGTTPELIIRDLDVVRRILNVDFAYFYPRGVGRNPKEEPVFLNLFHVDGDMWKLLRQRLTPTFTTAKLKNMFPLVVQCAEKLQTVGENIVNRGGDCDVRELMARFTTEFIGACGFGIQMDSINDEHSLFRALGKKMFQRSWKNYLVVALYDLFPEFRTIIQKSLRETEIMDAISEIVENIRKQRNYKPIGRNDFIDLLLELESKGKIVGESVEKRDANGKPEQVEMEMDLTCMVAQVFVFFAAGFETSSSATSFMLHQLAFHPEEQRKIQENIDQVLAKYDNKLCYDSISEMTALSNGFKEAMRLFPSLGTLHRVCAQKYTIPEMGITLDPGVKIIVPVQAIQVDGKYFENPTEFNPDRFNDTSADRHKFAYLPFGEGPRQCIGARLGEMQSLAGLAAVLHKFSVEPAANTKRHLEVNHGSNTVQSIKGGLPLKLRLRSKQSQVAA

> *S. litura* P450 6B6-like (GenBank: XP_022816422.1)

MFLLLLPLVLVLVYHYVTRNHDYWEKRNVKYEKPVPIFGTLYANLVAKKSITEISVEMYNKYPNEQAVGIYRGTTPELIIRDLDVVRRILNVDFAYFYPRGIGRNPKEEPVFLNLFHVDGDMWKLLRQRLTPTFTTAKLKNMFPLVVQCAEKLQTVGENIVSRGGDCDVRELMARFTTEFIGACGFGIQMDSINDEHSLFRALGKKMFQRSWKNYVAVALYDLFPEFRTIIQKSLRETEIMDAISEIVENIRKQRNYKPIGRNDFIDLLLELESKGKIVGESVEKRDANGKPEQVEMEMDLTCMVAQVFVFFAAGFETSSSATSFMLHQLAFHPEEQRKIQENIDQVLAKYDNKLCYDSISEMTALSNGFKEAMRLFPSLGTLHRVCAQKYTIPEMGITLDPGVKIIVPVQAIQVDGKYFDNPTEFNPDRFNDTSADRHKFAYLPFGEGPRQCIGARLGEMQSLAGLAAVLHKFSLEPAANTKRHLEVNHGSNTVQSIKGGLPLKLRLRSKQSQVAA

> *S. litura* P450 6B2-like (GenBank: XP_022829978.1)

MFLILVIIGLVALYFYGTRTFNYWKSRGIKHDKPVPIFGTNLKQFMQQASMCMMATEMYNKYPEEKVVGFYRGTDPELVIRDPEIIKRILTTDFQCFHSRGLTYHKTCVEPLLRNIFFADGDLWRLIRQRFTPAFSTGKLKAMFHIITDRAEKLQVITEEVVDREYYDARELMARYTTDFIGVCAFGIDMDSLSDENSHFRKLGKRIFERRFRDAVAGALKIMFPEVFKHMHFLAPELETNMKHLVQTVLKNRNYKPSGRNDFIDLMLELREKGKIIGESIEHKNPDGTPKIVDLEMDDLLMTAQAFVFFGAGFETSSTASSYTLHQLAFHPEYQKKVQEEVDTVLAKHNNKLTYDAIKEMKYLEMAFYESMRMYPSVGYLIRQCCVPKYTFPEIDLTIDDGLKVMIPIQAIHRDEKYFREPNKFDPERFSDGTKEDIKNFVYLPFGEGPRSCVGARLGQMQSMAGLAAVLQKYSVEPAACSRQEPLPDPAGIVSEGFVGGLPLKIRRRVK

> *S. exigua* CYP6AN4 (GenBank: ASO98001.1)

MFLLLLPLVLVLLYHYVTRNHDYWEKRNVKYEKPVPIFGTIYDNVVAKKSITEVSVELYNKYPNEPAVGIYRGTTPELVVRDLDVVRRILNVDFAYFYPRGIGRNPKEEPVFLNLFHVDGDKWKLLRQRLTPTFTTAKLKNMFPLVVQCAEKLQSVGEDIVNRGGDCDVRELMARFTTEFIGACGFGIQMDSINDEHSLFRALGRKIFQRTWKNYIAVALYDLFPEFRTIIQKSLRETKIMDSISEIVENIRKQRNYKPIGRNDFIDLLLELESKGKIVGESVEKRDANGKPEQVEMEMDLTCMVAQVFVFFAAGFETSSSATSFMLHQLAFHPKEQRKIQENIDQVLAKYDNKLCYDSISEMTALSNGFKEAMRLFPSLGTLHRVCAQKYTIPEMGITLDPGVKIIIPVQAIQVDEKYFENPTEFIPDRFNDTSADRHKFAYLPFGEGPRQCIGARLGEMQSLAGLAALLHKFSVEPAANTKRHLEVNHGSNTVQSIKGGLPLKLRLRSKESHVAA

> *C. includens* CYP6AN31 (GenBank: CAD0198413.1)

MITYLVIPIVLVLLYKYLTKNHDYWEKRNVKYEKPVPIFGTNYKNVTAQKSIIEISVEAYKKYPDEPVVGLLRGTTPELVVRDLDIARRILNVDFAYFYPRGTGRNPELEPLFYNLFHVDGDLWKLLRQRLTPTFTTAKLKNMFPLVVKCAEKLQHVGEDVVSRGGECDVRDLMARFTMEFIGACGFGIEMDTINNEQSLFKQLGNKIFRPYLRKHAINFLCDLFPDLSTILQKLLRESEIHDSITVILKNILEQRNYKPIGRNDFIDLLLELEGKGKIKGESVEKRKEDGTPEQVEMEMNLTYMAAQVFVFFAAGFETSSSATSFMLHQLAYHPEEQRKIQEEIDRVLSKYDNKVCYDSISEMTALSNGFKEALRLFPSLGALHRKCARRYTIPELNLTLDPGVKIIVPVQAIQNDEKYFDNPSQFLPDRFNDPNVGRHKYAYMPFGEGPRACIGARLGEMQSLAGLAAVLHKFSVEPGPRTQRYPEVNHGSNVIQSIKGGLPLKLSLRNKCNNN

> *C. includens* CYP6AN44 (GenBank: CAD0198411.1)

MITGSVQSSHERCGDNMIYLVILLVLVLLYKYTTKNHDYWEKRNVKYEKPVPIFGTDYKNVVGQKSLTEMSAELYNKYPNEPVVGLLRGTTPELVVRDLDVVRRILNVDFAYFYPRGLGRNPEVEPVFYNLFHVDGDLWKLLRQRLTPTFTTAKLKNMFPLVVKCAEKLQHVGEDVVSRGGECDVRDLMARFAMEFIGACGFGIQTDTINNDHSLFKELGNKIFRPYFRKYLMIACYDLFPDLRILLQKILHENDLIDAVTVIFKNIREQRNYKPIGRNDFIDLLLELEGKGKITGESVEKRDANGTPEQVEMEMNFTCMAAQVFLFFAAGFETSSSSTSFMLHQLAYHPEEQRKIQKEIDRVLNKYDNKLCYDSISEMTALSNGFKEAMRLFPSVGTLHRVCARRCTIPELNLTLDPGVKIIVPVQAIQNDEKYFDNPSQYLPDRFNDPNVGRHKYAYMPFGEGPRACIGARLGEMQSLAGLAAVLHKFSVEPGPRTLRYPEVNHGSNVIQSIKGGLPLKLTLRNKSNR

> *T. ni* P450 6B5-like (1) (GenBank: XP_026729651.1)

MIIYLVIPIVLVLLYKYVTRNHDYWEKRNVKYEKPISIFGTNYENVVAKKSIIEISIEIYNKYPNEPVVGLLRGTTPELVVRDLEIVRKILNVDFAHFYPRGTGRDPKLEPLFYNLFHVEGDLWKLLRQRLTPTFTTAKLKNMFPLVVKCAEKVQIVGEDIVNRGGECDVRDLMARFTMEFIGACGFGIEMDTINDEQSLFKQLGNKIFRPYFRKQAINALCDLFPDLQTVLQKLLRESDIIDAVTEILEKIREQRNYKPIGRNDFIDLLLELEGKGKIIGDSIEKNKADGTPEQVEMEMDLTYMAAQVFVFFAAGFETSSSATSFMLHQLAYHPEEQKKIQEEIDRVLSKYNNKLCYDSISEMTALSNGFKESLRIFPSLGALHRKCARRYTIPELNLTLDPGVKIIIPLQAIQNDEKYFDNPSQFMPDRFNDPNVGRHKYAYMPFGEGPRACIGARLGEMQSLAGLAAVLHKFSVEPAANTKRYPEVNHGSNIVQSIKGGLPLKLRLRDKSIS

> *T. ni* P450 6B5-like (2) (GenBank: XP_026729653.1)

MIYLLLPIVLLLLYKYFTKNHDYWEKRNVKYEKPIPIFGTNYENVIGKKSIIEKATEVYNKYPTEPVVGMFRGTTPELVVRDPEIARKILNVDFAYFYRRGLGKNPEIDPVYYNLFHVEGDLWKLLRQRLTPTFTTAKLKNMFPLIVNCAEKLRHVGEDIVNRGGECDVRDLMARFTVEFIGVCAFGIEMDTINNEHSLFKKLGVRIFKTDFRKYLVTALYDLFPDFRTIIQKLLSNSSSITAITEIFYNIRKQRNYKPIGRNDFIDLLLELEGKGKIIGDSIEKKNADGKPVQVEMEMNITYMAAQMFVFFAAGFETSSSATSFTLHQLAYHPEEQQKIQEEIDRVLSKYNNKLCYDSISEMTALSNGFKEAMRIFPSLGTLHRVCARKYTIPGLNVTLDPGVKIIIPTQGIQNDEKYFDNPSQFIPDRFNDPNVLRHKYAYLPFGGGPRACIGGRLGEMQSLAGLAAILHKFSVEPGPKTQRYPQVNHGSNLIQGVKGGIPLKLRLRNKSNSVAA
