## Supplementary file 2 for "Molecular and functional characterization of CYP6AN1 orthologs involved in capsaicin metabolism in two *Helicoverpa* species"

> Sequence of a in Fig. 1A

AAACTGGTTTGAACCATCCAGCTAGAACAACCTATATTTTAAAATTTCGCGAGTTTTTGCCAAATAGTGATCAAAATAAACATTTTTTTAATACGAAGAACTGAGGTCGAACAAACTACCAGCTATTTAGTAAAAAATATGGAATAGAGAAAAAGCACCAACAAAGAAGTGTTGATATACTCAAAAATAAGCTAAACAGAGTCGTTTGTTTCCAACAACTATCACGCCGCATTATAATCATCTAATATTGAACAATAAAACGTGTGTCTGTAATCGCTCACTATACGGCTGTGGCCCGACGGAACCCCAAGTAGGTACCGTTCACACAAACAACACAAGATAACGGGTCCCACCGCACTCTCCGCGGTTAACTCAGTCGCACGCACGTCCACAACATGATATTCCTTCTGCTACTGCCTATTGCACTCGTGCTCTTATACTACTACACCACCAGGAACCACGACTACTGGGAGAAACGCAATGTCAAGCACCAGAAACCAATACCCGTATTTGGAACGCTATACGACAATGTGTTTGCCAAGAAAAGCATAATAGAAATGGCCGCTGATTTATACCGCCAGTATCCCAATGAGAAAGTTGTG

> Sequence of b in Fig. 1A

AAACTGGTTTGAACCATCCAGCTAGAACAACCTATATTTTAAAATTTCGCGAGTTTTTGCCAAATAGTGATCAAAATAAACATTTTTTAATACGAAGAACTGAGGTATATTACTAAAATATAAAGACAATGTGATAAGAAAATGTCCTTATCACCTTGTCTAGATATTTTAAGTACCTACAGCTAGTGGAATATTGAGATAATAGCTGAAAGATAACAACGTACTTATAAAAATATACTCCAGTATCAAGCACTTACACAATTGTCGAACAAACTACCAGCTATTTAGTAAAAAATATGGAATAGAGAAAAAGCACCAACAAAGAAGTGTTGATATACTCAAAAATAAGCTAAACAGAGTCGTTTGTTTCCAACAACTATCACGCCGCATTATAATCATCTAATATTGAACAATAAAACGTGTGTCTGTAATCGCTCACTATACGGCTGTGGCCCGACGGAACCCCAAGTAGGTACCGTTCACACAAACAACACAAGATAACGGGTCCCACCGCACTCTCCGCGGTTAACTCAGTCGCACGCACGTCCACAACATGATATTCCTTCTGCTACTGCCTATTGCACTCGTGCTCTTATACTACTACACCACCAGGAACCACGACTACTGGGAGAAACGCAATGTCAAGCACCAGAAACCAATACCCGTATTTGGAACGCTATACGACAATGTGTTTGCCAAGAAAAGCATAATAGAAATGGCCGCTGATTTATACCGCCAGTATCCCAATGAGAAAGTTGTGGGTATGTTCCGAGGAAACACTCCGGAGCTCGTCATTCGAGACCTGGAAATAGTAAGGAAGATCCTGAATGTAGACTTCGCGTACTTCTACCCAAGAGGAATCGGCAGAAACCCT

> Sequence of c in Fig. 1A; *H. assulta CYP6AN1v1* transcript in Fig. 1B (GenBank: PQ470131.1)

AAACTGGTTTGAACCATCCAGCTAGAACAACCTATATTTTAAAATTTCGCGAGTTTTTGCCAAATAGTGATCAAAATAAACATTTTTTTAATACGAAGAACTGAGGTATATTACTAAAATATAAAGACAATGTGATAAGAAAATGTCCTTATCACCTTGTCTAGATATTTTAAGTACAGCTAGTGGAATATTGAGATAATAGCTGAAAGATAACAACGTATAAAAATATACTCCAGTATCAAGCACTTACACAATTGTCGAACAAACTACCAGCTATTTAGTAAAAAATATGGAATAGAGAAAAAGCACCAACAAAGAAGTGTTGATATACTCAAAAATAAGCTAAACAGAGTCGTTTGTTTCCAACAACTATCACGCCGCATTATAATCATCTAATATTGAACAATAAAACGTGTGTCTGTAATCGCTCACTATACGGCTGTGGCCCGACGGAACCCCAAGTAGGTACCGTTCACACAAACAACACAAGATAACGGGTCCCACCGCACTCTCCGCGGTTAACTCAGTCGCACGCACGTCCACAACATGATATTCCTTCTGCTACTGCCTATTGCACTCGTGCTCTTATACTACTACACCACCAGGAACCACGACTACTGGGAGAAACGCAATGTCAAGCACCAGAAACCAATACCCGTATTTGGAACGCTATACGACAATGTGTTTGCCAAGAAAAGCATAATAGAAATGGCCGCTGATTTATACCGCCAGTATCCCAATGAGAAAGTTGTGGGTATGTTCCGAGGAAACACTCCGGAGCTCGTCATTCGAGACCTGGAAATAGTAAGGAAGATCCTGAATGTAGACTTCGCGTACTTCTACCCAAGAGGAATCGGCAGAAACCCTACATATGAGCCTTTGTTTATGAATCTTTTCCATGCTGAAGGTGATTCGTGGAAGTTATTACGGCAGCGACTAACATCCACGTTCACGACTGCCAAGCTGAAAAAAATGTTTCCACTGGTAGTTGGGTGCGCTGAGAAACTGACACACGTTGGAGAAGACATCGTGAACAAAGGCGGAGACTGTGACGTGCGCGAACTGATGGCGCGCTTTACCACGGAGTTCATCGGCGCGTGCGGCTTCGGCATACAGATGGACACCATCAATGACGAGCACTCACTCTTCAGACATTTGGGCAGAAAAATGTTCAACAGATCTACGAAAATTCACATTCTCGTAGCACTATATGATCTATTTCCTGAGCTCGGGACATTCTTTCAAAAAACATTGAGGGAGACATTCCTACTGGATTCCATGACAGAGATAGTTCAAAATATTCGTAAACAACGTAACTTCAAGCCTATTGGCCGAAATGACTTCATTGATTTACTTTTAGAGCTGGAAGGAAAAGGTAAAATCACTGGGGAGTCGGTAGAGAAGAGAACTGCGGATGGAAAGCCAGTGCAAGTAGAAATGGAAATGGATCTGACATGCATGGTGGCACAGGTGTTCGTGTTCTTCGCAGCAGGGTTCGAAACTTCTTCATCGGCGACGAGTTTCATGTTACATCAGCTGGCATTCCACCCAGAAGAACAAAAGAAAATTCAGGAAGATATCGACAGGGTGCTTGCGAAGTACAACAACCAACTGTGCTATGACTCGATCTCTGAAATGAAAGCATTGAGTAACGGGTTCAAGGAGGCAATGAGAATGTTCCCCTCTCTCGGCACCTTACACAGAGTGTGTGCAAAGAAGTATAAAATCGAAGAGTTGGGCATCACAATCGATCCAGATGTGAAGATTATCATCCCACTGCAAGGTATACAAAATGACGGGAAATATTTTGAACATCCGGAGAAGTTCTTGCCGGATAGATTCAACGATGCGTCAGAAGAAAGAAATAAGTTTGCATACATGCCTTTCGGTGAAGGACCTCGGCAGTGCATCGGTGCTCGACTGGGCGAGATGCAGTCGCTGGCAGGACTTGCAGCAGTACTGCACAAGTTCAGTGTTGAGCCGACGACGACCACCAAGAGGTACCCAGACATCAACCACGCCAGCAACATTGTGCAATCAATCGAGGGAGGCTTGCCTCTCAAACTGAAACTTAGGAGCAAGGAAAATAATGTGGCTGCATAAGGAACACTTACTATAGATCTCCATTCAAAATAATATCTTGAACTGAAGTTAAATAGGTAGATAAATGAAGATGATAATTGAATAGCACCTAATATTGAATAAGGTGTTTTTAACATTCAAGGTTATTTATTAAAACCCAGTGATACCAAAATAAGCAATAACAATTACGTAGAACGAAATTCTATGGTCAATGGCCATAAAGTGAGTTTGAGTTTATAGGATTGAAATGAAGTATTACAGATGTTGTTTATATTTTATTCTTATAACTGTAATTTTAAGCACTAAAAATATATAAGATAATTTTTTATAAAACGCAAAAAAAAAAAAAAAAAA

> Sequence of d in Fig. 1A

ACAGCTAGTGGAATACTGAGATAATAGCTGAAAGATAACAACGTACTTATAAAAATATACTCCAGTATCAAGCACTTACACAATTGTCGAACAAACTACCAGCTATTTAGTAAAAAATATGGAATAGAGAAAAAGCACCAACAAAGAAGTGTTGATATACTCAAAAATAAGCTAAACAGAGTCGTTTGTTTCCAACAACTATCACGCCGCATTATAATCATCTAATATTGAACAATAAAACGTGTGTCTGTAATCGCTCACTATACGGCTGTGGCCCGACGGAACCCCAAGTAGGTACCGTTCACACAAACAACACAAGATAACGGGTCCCACCGCACTCTCCGCGGTTAACTCAGTCGCACGCACGTCCACAACATGATATTCCTTCTGCTACTGCCTATTGCACTCGTGCTCTTATACTACTACACCACCAGGAACCACGACTACTGGGAGAAACGCAATGTCAAGCACCAGAAACCAATACCCGTATTTGGAACGCTATACGACAATGTGTTTGCCAAGAAAAGCATAATAGAAATGGCCGCTGATTTATACCGCCAGTATCCCAATGAGAAAGTTGTG

> Sequence of e in Fig. 1A

ACACAATTGTGAGTAAACGGCACTGTTGAAATTCAACATAATTATCGTACCTTTTTGTAGTTTATCGATAGTTTCTTCAATGCCCTAAAATATTTATCTATTTTTCGGTCATTAGAACTAAAAAATCATTTAAAGAACTAAAAAATTTTGCAAAACCTTATATTAACACCCTTTTTCCTTTTGATATTTGAATAGGAAAATACCTACCTACTTATTTGTAGTTAGTTAAATACAAAGATAAATACTTCACCACATGCCAGAGACTTAATTGGTGTTGCACGTAGCAGGATATGTCTATATTAAATGTTGGTACCAATTATGCCAAAGACTCGTATGTCATTACATCCATTTGAAACATCCAATGCATCACAATGTATACAGCGCTGACAATTTATAGTACCTACATATGATGTAATTGTGATGCAATCTATAAACTTAAAACATATATCTTTTCATTACAGGTCGAACAAACTACCAGCTATTTAGTAAAAAATATGGAATAGAGAAAAAGCACCAACAAAGAAGTGTTGATATACTCAAAAATAAGCTAAACAGAGTCGTTTGTTTCCAACAACTATCACGCCGCATTATAATCATCTAATATTGAACAATAAAACGTGTGTCTGTAATCGCTCACTATACGGCTGTGGCCCGACGGAACCCCAAGTAGGTACCGTTCACACAAACAACACAAGATAACGGGTCCCACCGCACTCTCCGCGGTTAACTCAGTCGCACGCACGTCCACAACATGATATTCCTTCTGCTACTGCCTATTGCACTCGTGCTCTTATACTACTACACCACCAGGAGCCACGACTACTGGGAGAAACGCAATGTCAAGCACCAGAAACCAATACCCGTATTTGGAACGCTATACGACAATGTGTTTGCCAAGAAAAGCATAATAGAAATGGCCGCTGATTTATACCGCCAGTATCCCAATGAGAAAGTTGTGGGTATGTTCCGAGGAAACACTCCGGAGCTCGTCATTCGAGACCTGGAAATGGTAAGGAAGATCCTGAATGTAGACTTCGCGTACTTCTACCCAAGAGGAATCGGCAGAAACCCT

> Sequence of f in Fig. 1A; *H. assulta CYP6AN1v2* transcript in Fig. 1B (GenBank: PQ470132.1)

ACACAATTGTGAGTAAACGGCACTGTTGAAATTCAACATAATTATCGTACCTTTTTGTAGTTTATCGATAGTTTCTTCAATGCCCTAAAATATTTATCTATTTTTCGGTCATTAGAACTAAAAAATCATTTAAAGAACTAAAAAATTTTGCAAAACCTTATATTAACACCCTTTTTCCTTTTGATATTTGAATAGGAAAATACCTACCTACTTATTTGTAGTTAGTTAAATACAAAGATAAATACTTCACCACATGCCAGAGACTTAATTGGTGTTGCACGTAGCAGGATATGTCTATATTAAATGTCGAACAAACTACCAGCTATTTAGTAAAAAATATGGAATAGAGAAAAAGCACCAACAAAGAAGTGTTGATATACTCAAAAATAAGCTAAACAGAGTCGTTTGTTTCCAACAACTATCACGCCGCATTATAATCATCTAATATTGAACAATAAAACGTGTGTCTGTAATCGCTCACTATACGGCTGTGGCCCGACGGAACCCCAAGTAGGTACCGTTCACACAAACAACACAAGATAACGGGTCCCACCGCACTCTCCGCGGTTAACTCAGTCGCACGCACGTCCACAACATGATATTCCTTCTGCTACTGCCTATTGCACTCGTGCTCTTATACTACTACACCACCAGGAACCACGACTACTGGGAGAAACGCAATGTCAAGCACCAGAAACCAATACCCGTATTTGGAACGCTATACGACAATGTGTTTGCCAAGAAAAGCATAATAGAAATGGCCGCTGATTTATACCGCCAGTATCCCAATGAGAAAGTTGTGGGTATGTTCCGAGGAAACACTCCGGAGCTCGTCATTCGAGACCTGGAAATAGTAAGGAAGATCCTGAATGTAGACTTCGCGTACTTCTACCCAAGAGGAATCGGCAGAAACCCTACATATGAGCCTTTGTTTATGAATCTTTTCCATGCTGAAGGTGATTCGTGGAAGTTATTACGGCAGCGACTGACATCCACGTTCACGACTGCCAAGCTGAAAAAAATGTTTCCACTGGTAGTTGGGTGCGCTGAGAAACTGACACACGTTGGAGAAGACATCGTGAACAAAGGCGGAGACTGTGACGTGCGCGAACTGATGGCGCGCTTTACCACGGAGTTCATCGGCGCGTGCGGCTTCGGCATACAGATGGACACCATCAATGACGAGCACTCACTCTTCAGAGATTTGGGCAGAAAAATGTTCAACAGATCTACGAAAATTCACATTCTCGTAGCACTATATGATCTATTTCCTGAGCTCGGGACATTCTTTCAAAAAACATTGAGGGAGACATTCCTACTGGATTCCATGACAGAGATAGTTCAAAATATTCGTAAACAACGTAACTTCAAGCCTATTGGCCGAAATGACTTCATTGATTTACTTTTAGAGCTGGAAGGAAAAGGTAAAATCACTGGGGAGTCGGTAGAGAAGAGAACTGCGGATGGAAAGCCAGTGCAAGTAGAAATGGGAATGGATCTGACATGCATGGTGGCACAGGTGTTCGTGTTCTTCGCAGCAGGGTTCGAAACTTCTTCATCGGCGACGAGTTTCATGTTACATCAGCTGGCATTCCACCCAGAAGAACAAAAGAAAATTCAGGAAGATATCGACAGGGTGCTTGCGAAGTACAACAACCAACTGTGCTATGACTCGATCTCTGAAATGAAAGCATTGAGTAACGGGTTCAAGGAGGCCATGAGAATGTTCCCCTCTCTCGGCACCTTACACAGAGTGTGTGCAAAGAAGTATAAAATCGAAGAGTTGGGCATCACAATCGATCCAGATGTGAAGATTATCATCCCACTGCAAGGTATACAAAATGACGAGAAATATTTTGAACATCCGGAGAAGTTCTTGCCGGATAGATTCAACGATGCGTCAGAAGAAAGAAATAAGTTTGCATACATGCCTTTCGGTGAAGGACCTCGGCAGTGCATCGGTGCTCGACTGGGCGAGATGCAGTCGCTGGCAGGACTTGCAGCAGTACTGCACAAGTTCAGCGTTGAGCCGACGACGACCACCAAGAGGTACCCAGACATCAACCACGCCAGCAACATTGTGCAATCAATCGAGGGAGGCTTGCCTCTCAAACTGAAACTTAGGAGCAAGGAAAATAATGTGGCTGCATAAGGAACACTTACTATAGATCTCCATTCAAAATAATATCTTGAACTGAAGTTAAATAGGTAGATAAACGAAGATGATAATTGAATAGCACCTAATATTGAATAAGGTGTTTTTAACATTCAAGGTTATTTATTAAAACCCAGTGATACCAAAATAAGCAATAACAATTATGTAGAACGAAATTCTATGGTCAATGGCCATAAAGTGACTTTGAGTTTATAGGATTGAAATGAAGTATTACAGATGTTGTTTATATTTTATTCTTATAACTGTAATTTTAAGCACTAAAAATATATAAGATAATTTTTTATAAAACGCAAAAAAAAAAAAAAAAAA

> Sequence of g in Fig. 1A; *H. assulta CYP6AN1v3* transcript in Fig. 1B (GenBank: PQ470133.1)

ACACAATTGTGAGTAAACAGCACTGTTGAAATTCAACATAATTATCGTCGAACAAACTACCAGCTATTTAGTAAAAAATATGGAATAGAGAAAAAGCACCAACAAAGAAGTGTTGATATACTCAAAAATAAGCTAAACAGAGTCGTTTGTTTCCAACAACTATCACGCCGCATTATAATCATCTAATATTGAACAATAAAACGTGTGTCTGTAATCGCTCACTATACGGCTGTGGCCCGACGGAACCCCAAGTAGGTACCGTTCACACAAACAACACAAGATAACGGGTCCCACCGCACTCTCCGCGGTTAACTCAGTCGCACGCACGTCCACAACATGATATTCCTTCTGCTACTGCCTATTGCACTCGTGCTCTTATACTACTACACCACCAGGAACCACGACTACTGGGAGAAACGCAATGTCAAGCACCAGAAACCAATACCCGTATTTGGAACGCTATACGACAATGTGTTTGCCAAGAAAAGCATAATAGAAATGGCCGCTGATTTATACCGCCAGTATCCCAATGAGAAAGTTGTGGGTATGTTCCGAGGAAACACTCCGGAGCTCGTCATTCGAGACCTGGAAATAGTAAGGAAGATCCTGAATGTAGACTTCGCGTACTTCTACCCAAGAGGAATCGGCAGAAACCCTACATATGAGCCTTTGTTTATGAATCTTTTCCATGCCGAAGGTGATTCGTGGAAGTTATTACGGCAGCGACTAACATCCACGTTCACGACTGCCAAGCTGAAAAAAATGTTTCCACTGGTAGTTGGGTGCGCTGAGAAACTGACACACGTTGGAGAAGACATCGTGAACAAAGGCGGAGACTGTGACGTGCGCGAACTGATGGCGCGCTTTACCACGGAGTTCATCGGCGCGTGCGGCTTCGGCATACAGATGGACACCATCAATGACGAGCACTCACTCTTCAGACATTTGGGCAGAAAAATGTTCAACAGATCTACGAAAATTCACATTCTCGTAGCACTATATGATCTATTTCCTGAGCTCGGGACATTCTTTCAAAAAACATTGAGGGAGACATTCCTACTGGATTCCATGACAGAGATAGTTCAAAATATTCGTAAACAACGTAACTTCAAGCCTATTGGCCGAAATGACTTCATTGATTTACTTTTAGAGCTGGAAGGAAAAGGTAAAATCACTGGGGAGTCGGTAGAGAAGAGAACTGCGGATGGAAAGCCAGTGCAAGTAGAAATGGAAATGGATCTGACATGCATGGTGGCACAGGTGTTCGTGTTCTTCGCAGCAGGGTTCGAAACTTCTTCATCGGCGACGAGTTTCATGTTACATCAGCTGGCATTCCACCCAGAAGAACAAAAGAAAATTCAGGAAGATATCGACAGGGTGCTTGCGAAGTACAACAACCAACTGTGCTATGACTCGATCTCTGAAATGAAAGCATCGAGTAACGGGTTCAAGGAGGCAATGAGAATGTTCCCCTCTCTCGGCACCTTACACAGAGTGTGTGCAAAGAAGTATAAAATCGAAGAGTTGGGCATCACAATCGATCCAGATGTGAAGATTATCATCCCACTGCAAGGTATACAAAATGACGAGAAATATTTTGAACATCCGGAGAAGTTCTTGCCGGATAGATTCAACGATGCGTCAGAAGAAAGAAATAAGTTTGCATACATGCCTTTCGGTGAAGGACCTCGGCAGTGCATCGGTGCTCGACTGGGCGAGATGCAGTCGCTGGCAGGACTTGCAGCAGTACTGCACAAGTTCAGCGTTGAGCCGACGACGACCACCAAGAGGTACCCAGACATCAACCACGCCAGCAACATTGTGCAATCAATCGAGGGAGGCTTGCCTCTCAAACTGAAACTTAGGAGCAAGGAAAATAATGTGGCTGCATAAGGAACACTTACTATAGATCTCCATTCAAAATAATATCTTGAACTGAAGTTAAATAGGTAGATAAATGAAGATGATAATTGAATAGCACCTAATATTGAATAAGGTGTTTTTAACATTCAAGGTTATTTATTAAAACCCAGTGATACCAAAATAAGCAATAACAATTACGTAGAACGAAATTCTATGGTCAATGGCCATAAAGTGAGTTTGAGTTTATAGGATTGAAATGAAGTATTACAGATGTTGTTTATATTTTATTCTTATAACTGTAATTTTAAGCACTAAAAATATATAAGATAATTTTTTATAAAACGCAAAAAAAAAAAAAAAAAA

>*H. armigera CYP6AN1v2* transcript in Fig. 1B (GenBank: PQ470130.1)

TCTCAAACTGGTTTGAACCATCCAGCTAGAACAACCTATGCTAGAATTTCGCGAGTTTTTGCCAAATTGTGATAAAAATAAACATTTTTTAACACGGAACAACTGAGGTCGAACAAACTACCAGCTATTCAGTAAAAATCTGAAATAGAGGAAAGGTACCAACAAAGAAGTGTTGATAGACCGCAAAAATAAGCTAAACAGAGTCGTTTGTCTCCAACAACTATCGCGCCGCATTATACCTAATCATCTAATATTGAACAATAAAACGTGTGTCTGTAATCGCCCACTATACGGCTGTGGCCCGACGGAACCCCAAGTAGGTACCGTTCACACAAACAACACAAGATAACGGGTCCCACCGCACACTTCGCGGTTAACTCAGTCGCACGTCTTTAACCACGAACACAACATGATATTCCTGCTGCTACTGCCTATTGCACTCGTGCTCTTATACAAATACACCACCAGGAACCACGACTACTGGGAGAAACGCAACATCAAGCACGAGAAACCAATACCCGTATTTGGAACGCTATACGACAATGTGTTTGCCAAGAAAAGCATAATAGAAGTGGCCGCTGACTTATACCGCCAGTATCCCAATGAGAAAGTTGTGGGCATGTTCCGGGGAAACACTCCGGAGCTCGTCATTCGAGACCTGGAAATAGTGAGGAAGATCCTGAATGTAGACTTCGCGTACTTCTACCCAAGAGGAATCGGCAGGAACCCTACATATGAGCCTCTGTTTATGAATCTTTTCCATGCTGAGGGTGACTCGTGGAAGTTATTACGGCAGCGACTAACATCCACGTTCACGACTGCCAAGCTGAAAAAAATGTTTCCACTGGTAGTTGGGTGCGCTGAGAAACTGACACACGTTGGAGAAGACATCGTGAACAAAGGCGGAGACTGTGACGTGCGCGAGCTGATGGCGCGCTTTACCACGGAGTTCATCGGCGCGTGCGGCTTCGGCATACAGATGGACACCATCAATGACGAGCACTCACTCTTCAGAGATTTGGGCAGAAAAATGTTCAACAGATCGACGAAAATTCATTTCCTCGTAGCACTGTATGATCTATTTCCTGAGTTTGGGACTTTCTTTCAAAAAACATTGAAGGAGACATTCCTACTGGATTCCATGACAGAGATAGTTCAAAATATTCGTAAACAACGTAACTTCAAGCCCATCGGCCGAAATGACTTCATTGATTTACTTTTAGAGCTGGAAGAAAAAGGTAAAATCACAGGGGAGTCGGTGGAGAAGAGAACTGCGGATGGAAAACCGGTGCAAGTAGAAATGGAAATGGATCTGACATGCATGGTGGCGCAAGTGTTTGTGTTCTTCGCAGCAGGGTTCGAAACTTCCTCATCGGCAACGAGTTTCATGTTACATCAGTTGGCATTCCACCCAGAAGAACAAAAGAAAATTCAGGAAGATATCGATAGGGTTCTTGCGAAGTACAACAATCAACTGTGCTATGACTCGATCTCTGAAATGAAAGCATTGAGTAACGGGTTCAAAGAGGCAATGAGAATGTTCCCCTCTCTCGGCACCTTACACAGAGTGTGTGCAAAGAAGTACAAGATCGAAGAGTTAGGCATCACAATCGATCCTGATGTGAAGATTATCATCCCACTGCAAGGCATACAAAATGACGAGAAATATTTTGAACATCCGGAGAAGTTCATGCCGGATAGATTCAACGATACATCGGAAGAAAGAAATAAGTTTGCATACATGCCTTTCGGTGAAGGACCTCGGCAGTGCATCGGTGCTCGACTGGGTGAGATGCAGTCGCTGGCAGGACTTGCAGCAGTACTGCACAAGTTCAGCGTGGAGCCGACGACGACCACCAAGAGGTACCCAGACATCAACCACGGCAGTAACATTGTGCAATCCATCGAGGGAGGCCTACCACTGAAACTGAAACTTAGGAGCAAGGAAAATAATGTTGCTGCATAAGGAAGACTTACTATAGATCGCCATTCAAATAAATATTTTGAAGTTAAATAAGTACATAAATGAAGATGATAATTGAACAGCACCTAATATTGAACAAGGTGTTTTTAATATTCAAGGTTATTTATAGGCACCCAGTGATACCAAAATAAGCAATAACAATTACGTAGACCGAAAACGTATGGTCATTAAACTTGCTATGACCATTGATTTTATAGGATTACAATGAAGTATTACAGATGTTGGTTCTATTATTCTTATAACTGTAATTTTAAGCATCTAAAATATATAAGATAGAATTTTAAAAAAAAAAAAAAAAAAA

>Sequence spanning the *H. assulta CYP6AN1* gene and flanking regions

GGTACTTTCCATTACCTCACTCTAACAGTCTGATGAGACGACAATCTGACACTAGCGGAGAGGGTACGACAACTCATGCTTACTCGCAAACTGATGAGTTTGTTTGAACTAAAAAAAGACAATGTTATACCTCTATTATTTTCAGATTTCGAAACAAATAGCCAGTGACTAGCAATCAAAATTGTTTTTTCGTATAATTTGGTTTTTGTATAATTATTTGGAATTGGTCACTAAACTTGAGAACCTACTCTGTTTTGGGACCGGTTAAAACAGTTTTACCCCAGTATAGTTACCCAAAATACCGTTAGCAAATAGTGCATGCGCCCTGCATTTGTACAAAGTCAAAGTCAAATCATATATTTCATTTAGGCACTTACAAGCACTTATGAACGTCAAAAATTATAAATAAGTTTGATTCTAGTTGCACATTTCAAAAGTAGCTTCCTATGGAGAAGAACGGGCAAGAAACTCCATGGTTACTCTTTTTAAAACATAATAGGTTACAGTTTGTATGTATTGATACATACGATAATAACGTAAAAAAGAAGTGCTTTTATTATTAAATTCAATTTTCATTGATGTTATTAAAAAACACTTTGACATCAACACATTAGTTTAAATTAATTATTAATTCTAATCAAGGTTACTCATTCAACTTAATTACCTACCCACTTATTTAGTAACAATGTCCTATTGTCTATCAGGAAAAATATACCTATATATCTTCATAAATGCTCCAATCTTTTAAATCTAACCAATCAATTAAAAACATAAATACACATCCGTCCATAGTCTGACATTTGTCATATTACTCTAAAATTTTCGTCTACGAGCGACCGAACAAAGTCCAAAGTCTATAATGTACCGAGATATATAATACTGAGTTGTGAGGTTCTCAACTCAAACTGGTTTGAACCATCCAGCTAGAACAACCTATATTTTAAAATTTCGCGAGTTTTTGCCAAATAGTGATCAAAATAAACATTTTTTTAATACGAAGAACTGAGGTATGTATTATTAGATAAAGTGTTTTCTTAACCTTTGATAGTGTAAATAAGTTATTATCAAGCGTAAGGAGATTGTGTTAAAAATAGACAACTTGAGTCCAAGGGTCGATTTTTAGCTATATTTGGTGTGTGAAGAAAGTAAATTTCTTTAGTGAACGGTTAAAGTGACATTTATGACCAAAGTCTAGATCAATCATGATAGTATCTTGATTTATTAGTGTTTATTATCAACTTATCGGTGTTTTATTGGGTGTTAAGCCTTTGTTTATCGTATCAAAACATGTTCAAAATCTCGTCTATTTATGAAGTAGTTAATTTACTTGCTACATCATTTACATTTCCCGACAATTTATGCAAATTAAGCTAAACTTTATTTTTATTGTATCGTGGGTACCTACATTGTAATATCTTATTCTTGTCTCACCTTTGATAACTTTTTACTTTTAGGTATATTACTAAAATATAAAGACAATGTGATAAGAAAATGTCCTTATCACCTTGTCTAGATATTTTAAGTACAGCTAGTGGAATATTGAGATAATAGCTGAAAGATAACAACGTACTTATAAAAATATACTCCAGTATCAAGCACTTACACAATTGTGAGTAAACGGCACTGTTGAAATTCAACATAATTATCGTACCTTTTTGTAGTTTATCGATAGTTTCTTCAATGCCCTAAAATATTTATCTATTTTTCGGTCATTAGAACTAAAAAATCATTTAAAGAACTAAAAAATTTTGCAAAACCTTATATTAACACCCTTTTTCCTTTTGATATTTGAATAGGAAAATACCTACCTACTTATTTGTAGTTAGTTAAATACAAAGATAAATACTTCACCACATGCCAGAGACTTAATTGGTGTTGCACGTAGCAGGATATGTCTATATTAAATGTTGGTACCAATTATGCCAAAGACTCGTATGTCATTACATCCATTTGAAACATCCAATGCATCACAATGTATACGGCGCTGACAATTTATAGTACCTACATATGATGTAATTATGATGCAATCTATAAACTTAAAACATATATTTTTTCATTACAGGTCGAACAAACTACCAGCTATTTAGTAAAAAATATGGAATAGAGAAAAAGCACCAACAAAGAAGTGTTGATATACTCAAAAATAAGCTAAACAGAGTCGTTTGTTTCCAACAACTATCACGCCGCATTATAATCATCTAATATTGAACAATAAAACGTGTGTCTGTAATCGCTCACTATACGGCTGTGGCCCGACGGAACCCCAAGTAGGTACCGTTCACACAAACAACACAAGATAACGAGTCCCACCGCACTCTCCGCGGTTAACTCAGTCGCACGCACGTCCACAACATGATATTCCTTCTGCTACTGCCTATTGCACTCGTGCTCTTATACTACTACACCACCAGGAACCACGACTACTGGGAGAAACGCAATGTCAAGCACCAGAAACCAATACCCGTATTTGGAACGCTATACGACAATGTGTTTGCCAAGAAAAGCATAATAGAAATGGCCGCTGATTTATACCGCCAGTATCCCAATGAGAAAGTTGTGGGTATGTTCCGAGGAAACACTCCGGAGCTCGTCATTCGAGACCTGGAAATAGTAAGGAAGATCCTGAATGTAGACTTCGCGTACTTCTACCCAAGAGGAATCGGCAGAAACCCTACATATGAGCCTTTGTTTATGAATCTTTTCCATGCTGAAGGTGATTCGTGGAAGTTATTACGGCAGCGACTAACATCCACGTTCACGACTGCCAAGCTGAAAAAAATGTTTCCACTGGTAGTTGGGTGCGCTGAGAAACTGACACACGTTGGAGAAGACATCGTGAAACAAAGGCGGAGACTGTGACGTGCGCGAACTGATGGCGCGCTTTACCACGGAGTTCATCGGCGCGTGCGGCTTCGGCATACAGATGGACACCATCAATGACGAGCACTCACTCTTCAGAGATTTGGGCAGAAAAATGTTCAACAGATCTACGAAAATTCACATTCTCGTAGCACTATATGATCTATTTCCTGAGCTCGGGACATTCTTTCAAAAAACATTGAGGGAGACATTCCTACTGGATTCCATGACAGAGATAGTTCAAAATATTCGTAAACAACGTAACTTCAAGCCTATTGGCCGAAATGACTTCATTGATTTACTTTTAGAGCTGGAAGGAAAAGGTAAAATCACTGGGGAGTCGGTAGAGAAGAGAACTGCGGATGGAAAGCCAGTGCAAGTAGAAATGGAAATGGATCTGACATGCATGGTGGCACAGGTGTTCGTGTTCTTCGCAGCAGGGTTCGAAACTTCTTCATCGGCGACGAGTTTCATGTTACATCAGCTGGCATTCCACCCAGAAGAACAAAAGAAAATTCAGGAAGATATCGACAGGGTGCTTGCGAAGTACAACAACCAACTGTGCTATGACTCGATCTCTGAAATGAAAGCATTGAGTAACGGGTTCAAGGAGGCCATGAGAATGTTCCCCTCTCTCGGCACCTTACACAGAGTGTGTGCAAAGAAGTATAAAATCGAAGAGTTGGGCATCACAATCGATCCAGATGTGAAGATTATCATCCCACTGCAAGGTATACAAAATGACGAGAAATATTTTGAACATCCGGAGAAGTTCTTGCCGGATAGATTCAACGATGCGTCAGAAGAAAGAAATAAGTTTGCATACATGCCTTTCGGTGAAGGACCTCGGCAGTGCATCGGTGAGTAGCTAAAACATTGACTTATTAACTAGACACTTATTGTTCTAGACTGCGTTGTTCAGTTACTCCTTTCAAATTATAAAGTCTAATTTACACTTGCGAGTTAATAGCCCACTAGCGCAAAATGATAATTCAAACAAGCCCAGTTCATTCACAGTTTTATATGAACGGCAAACAAATAAGAATTACATTTCTAACCTTACATAATGTTTGTCTTCCAGGTGCTCGACTGGGCGAGATGCAGTCGCTGGCAGGACTTGCAGCAGTACTGCACAAGTTCAGCGTTGAGCCGACGACGACCACCAAGAGGTACCCAGACATCAACCACGCCAGCAACATTGTGCAATCAATCGAGGGAGGCTTGCCTCTCAAACTGAAACTTAGGAGCAAGGAAAATAATGTGGCTGCATAAGGAACACTTACTATAGATCTCCATTCAAAATAATATCTTGAACTGAAGTTAAATAGGTAGATAAATGAAGATGATAATTGAATAGCACCTAATATTGAATAAGGTGTTTTTAACATTCAAGGTTATTTATTAAAACCCAGTGATACCAAAATAAGCAATAACAATTATGTAGAACGAAATTCTATGGTCAATGGCCATAAAGTGACTTTGAGTTTATAGGATTGAAATGAAGTATTACAGATGTTGTTTATATTTTATTCTTATAACTGTAATTTTAAGCACTAAAAATATATAAGATAATTTTTTATAAAACGCACGAGTTTCTTTTATTTGCATCTATTTCCATAGTGAGTCATCCGGACTGGTTTAGCCAACAGATAGGCATTTTAATAACTTGCAATTATCTTTGATATTTGAGTGTTTCTCAAGAATTATATATGTCACCGGAAAATAACAGTTTAAAGACATACAATCGTTATTACCTTCTGTTTAAATAACATCAATGTTGATGTTAATCAATTCAGGGCCGTGAAATATAAATTTGCTGATATCTGCCATTTCTGAATCCTTTTGTCTTTATCATTTATATATAGTTCCAGTTGTTCCGAGGAAACTGTTCCCGATCCAAGATAAGTAACAGTCTCTAACATTATGACAACACTATGCTATGCACCAAACGCATTCCCGATTAAAGAGTTAGTGAAAATGTAGAATCATAGAAAAAATATTTTCCTGCCGTTATTATTACATATTTAGTTTATAAATATCTGTATCGGTTTCCCGTTTAACGTTAATAAATAAACAGACTGATATGATTTAGTACCCCTAACACTAAACGTTGCAATTCACACCGTATTTATGCAGATACTCAATGAACGATAACAAGTTACAACGTTCACTACACTGTTGATTCAGTCTCGCGCGCCTAGCCGCGGCATCAACATGCTTCTCCTGCTTATACCAATTGTGCTCGTGCTCCTATACTACTACACCACCAGGAACCACGACTACTGGGAGAAACGCAACGTCAAGCACGAGAAACCAATACCCGTGTTTGGAACGCTGTACAACAATGTATTTGCTAAGAAAAGCATCACCGAAATATCCGTGGAGCTATATCGACAGTATCCTAATGAGAAAGTTGTTGGAATATACAGAGGAACCACACCAGAACTCATCATCCGGGATCTGGATATAGCAAGGAAGATCCTGAATGTAGACTTCGCATATTTCTACCCCAGAGGACTGGGCAGGAATCCTAAATATGAACCCGCGTTTCTGAATATCTTCCATGTTGACGGTGACACGTGGAAATTGTTACGACAACGCCTAACATCAGCGTTCACGACCGCCAAACTGAAAAGCATGTTCCCACTAGTAGTGGCGTGCGCTGAAAAACTGCAAAATGTGGGCGAAGACATCGTGAGCCGTGGTAGAGACTGTGACGTGCGCGAGCTGATGGCGCGCTTTACCACCGAGTTCATCGGCGCGTGCGGCTTTGGTATTCAAATGGACACCATCAACAACGAGCACTCTCTCTTCAGAGATTTGGGCAGAAAAATGTTCAACAGATCGTGGAGAAATTTCGTCCTCGTACCACTCTACGATTTGTTTCCCGACTTCAGAACACTTTTTCAAAAAATATTGCACGAGCCGTTCTTACTGGATTCCATGGCCGAGATTGTTCAAAATATTCGCAAACAACGTAACTTCAAACCCATTGGCAGAAATGACTTCATTGATTTACTTTTAGAAGTAGAAGCCAAAGGCAAGATGACAGGTGAGTCCGTGGAGAAGAGAACTGCGGATGGAAAGCCAGTGCAAGTAGAAATGGACATGGATCTGACATGCATGGTGGCACAAGTGCTCGTGTTCTTCGCAGCAGGTTTCGAAACTTCCTCATCGGCGACGAGTTTCCTGTTACATCAGCTGGCATTCCATCCAGAAGAACAAAAGAAAATTCAGGAAGATATTGACAGGGTGCTTGCGAAGTACAACAACCAACTGTGCTATGACCCGATCTCTGAAATGAAAGCATTGAGTAACGGGTTCAAGGAAGCAATGAGAATGTTCCCCTCTCTTGGCACCCTACACAGAGTGTGTGCAAAGAAGTATAAAATCGAAGAGTTGGACATCACAATCGATCCAGATGTGAAGATTATCATTCCAGTGGAAGGTATACAAAATGACGAAAAGTATTTCGAAAACCCAACTCAGTTTAAACCGGATAGGTTCAACGATCCGTCTGAGAAGAGACATAAGTTTGCATACATACCCTTCGGTGAAGGACCTCGGCAATGTATCGGTGAGTAGTTTTTATATTGTTTTTCTCACAAAGACCTTTGTTTCGTTAAGTCATTTATAGTAATCTAACCTTGATATTGAAGCTGATAACTTTCACCAAAATACTACCTTCAATAAATGGTACGATGGCCAATGCTTTTCATATTATCGGCTCTAGATTAAAGCAATAAAATTATTCATCTGGAACCGAAATTTTCTGTCAAGTGGAAAATCCACGCGGATAACTTCAACGTGATAAGACATCAACCCAACTGCAATATTCCTGGATACCTATGTCCTTCGATATTTGAGGACCTTTTTTTCGGGGCTACAATTATGATTTATAAGGCCTTAGTCTGTTACAGTCTTCCATTTCTTTTTTGTCCACCATTAACACGAAGGTTCTCTATCGTCAATGATTTTCTCATCAGGTGCTCGGTTGGGCGAGATGCAGTCGCTAGCAGGACTTGCAGCAGTACTGCATAAGTTCAGCGTGGAGCCGGCAGCGACCACCAAGAGGTACCCCGAGGTGAACCACGGTAGTAATACGGTGCAGTCAATCAAGGGAGGACTGCCACTGAAGTTGACTCTGAGAAATAAATAGTTTTTCATAAAACTAACACTGTCCACGATGCAGACACGAGAGTCAAACAATCGGTATGCATCAAATATGACCCATGGGATAGGGAAAATTATTAAGAGCTCACAATCCAGACTTGAATAGAATACTTGAATAAAATACTTCTATCGAAAATAGTAAGAGCTCCGAAAACCATATAACCATATAAAAGGATTTAGCCCTTTATCATTAGCTTTTAATCGTACCTAAATAGATAATTATTATTGTAAGAATAAATCAATAAATATTAAGATAA

>Sequence spanning the *H. armigera CYP6AN1v2* gene and flanking regions

GTTACTTTCCATTACCTCACTCTATTTAAGGTGAGATGACAATCTGACACTAGCGGAGAGAGTAAGACAACTCATGCTTACTCGCAAACTGATGAGTTTGTTTGAACGCCCTAATCTAAGAAAAGACAGCACCGACATGAAAAAAAACAAATAATACTTACAATGTTGTAATATTTTCAGATAGCCTATTTATCGAGGTTTGCCTGTCATCAAAGGTTAATACAAATAGCCAGTGACTAGCAATCAACATTATTTTTTGTACCTATCGGTAATTTGTTTTTGTATAATTATTTGGAATTGGTCACTAAACTTGAGAACCTACTTTGTATTGGGATAGGCTAAAACAATTTTCAGTATAGTTACCCAAAATACTGTTAGCAAATAGTGCATGCGCCCTGCATTTGTTCAAAGTCAAAGTCAAATCCTTTATTTCATTCAGGCCTTTACAAGCACTTATGAACGTCAAAAATTAAAATAAGTTTGATTCTAGTTGCACATTCTAAAAGTAGCTTCCTATGGAGAAGCACGGGCAAGAAACTCCATGGATCATAATGTAAAAAAGAAGTGCTTACAATATTTTATGGGTTGAATCTGAGAAAGGTATACAATGCAAGTTCAACTAGGAAGTTATATGAGAAAATTATTCAAATCAATTTTCATTGATATTCTTAAAAAAATTATTCGACATCAACACATTAGTTTAAATTAATTATTAATTCTAACCAAGGTTAGTCATTCAACTTAATTACCTACCCACTTATTTAGCAACAATGTTCTATTATCTACCAGGAAAAATATACATACCTTCATAAGTACTCCAATCTTTAAAATCTAACCAATCAATTAAAAACTAATACACATTCATCAATAGTCTGACTTTTGTCATAATACTCTAAAATTTTTGTCTACGAGCGACCGAACAAAGTCCAAAGTCTATATTGGACCGAGATATATAATACTGAGTTATGAGGCTCTCAACTCAAACTGGTTTGAACCATCCAGCTAGAACAACCTATGCTAGAATTTCGCGAGTTTTTGCCAAATAGTGATCGAAATAAACATTTTTTAACACGGAAGAACTGAGGTATGTATTGTTTGATAAAGTGTTTTCTTAACCTTTGGTAGTGTAAATAAGTTATTATCAAGTGTAAGGAGATTGTGTTAAAAATAGACAACTTGAGTCCAAAGGTCGATTTTTAGCTTTATTTGGTGTCTGAAGAAAGTAACTTTCTTTAGTGAACGGGTAAAGTGGCATTTATGATCAAAGTCTAGATCAATCATGATAGTATCATGATTTATTTGTGTTTATTAACTTATCGGTGTTTTATTGGGTGTTAAGCCTTTGTGTATCGTATCAAAACATGTTCAAAATCTCATCTATTTTTGAAGTAGGTAATTATTTGCTACATCATTTACATTTTCCGATTTTTTTTTGCAAATTATGCTAAACTTTATTTTTATTGTATCGTGGGTACCTACCTTCATTGTAATATCTTATTCTTGTCTCACCTTTGATAACTATTTACTATTTTATTACTAAGGAAGGTGATAAGGACATGTTCTTATCACCTTGTCTAGGCATACAGCTGGTAGAATATTGAGATAATAGTTGAAAGATAACAACGTGTAAAAATATATTCCAGTATCAACCAGTTACACAACTTGTGTGAGTAAACAGCACTGTTGAAATTCAATACAATTATCGTATCTTTTTGTAGTTAATCGATAGTTTCTTCAATGCCCTCAAATAATTTTCGATCATTAGAACAAAAAATTCAATTCAGTTTTTCGAAAAGATTATTTTAACTCAATAGAAGTAAGTCCTTTTGTAGTAATACCTTTTTTCCTTTTGGCATTTGAGTAGGAAAATTGATTAAAAATCACATCTTATCATTGATATTATTTATTTATAGAGTAGGCACCTTCATACTTATTTTTAGTTAGTTAAATACAAAGATAGATGCTTCATCACATGCCAGAGACTTAATTGGTGATGCACGTAGCAGGATATGTCTATAATATTAAATGTTGGTGCCAATTATGCCAAAGACTCGTATGTCATTACATCCATTTGAAACATACAATTTAGCACAATGTGTATAGAGCTAGTATAGTACCTACCTATGATGAGATTGTGATGCAATCTGTGAACTTATTTTTTTTTTTCATTACAGGTTGAACAACTTACCAGCTATTTAGTAAAAATCTGATATAGAGGAAAGGTACCAACAACTGTTGATAAACCGCAAAAATAAGCTAAACAGAGTCGGTTGTTTCAAACAACTATCGCGCCGCATTATAATCATCTAATATAGAACAATAAAACGTGTGTCTGTAATCGCCCACTATACGGCTGTGGCCCGACGGAACCCCAAGTAGGTACCGTTCACACAAACAACACAAGATAACGGGTCCCACCGCACACTCCGCGGTTAACTCAGTCGCACGTCTTTAACCACGAACACAACATGATATTCCTGCTGCTACTGCCTATTGCACTCGTGCTCTTATACAAATACACCACCAGGAACCATGACTACTGGGAGAAACGCAACGTCAAGCACGAGAAACCAATACCCGTATTTGGAACGCTATACGACAATGTGTTTGCCAAGAAAAGCATAATAGAAGTGGCCGCTGATTTATACCGCCAGTATCCCAATGAGAAAGTTGTGGGCATGTTCCGGGGAAACACTCCGGAGCTCGTCATTCGAGACCTGGAAATAGTAAGGAAGATCCTGAATGTAGACTTCGCGTACTTCTACCCAAGAGGAATCGGCAGAAACCCTAAATATGAGCCTCTGTTTATGAATCTTTTCCATGCTGAAGGGGATTCGTGGAAGTTATTACGGCAGCGACTAACATCCACGTTCACGACTGCCAAGCTGAAAAAAATGTTCCCACTGGTAGTTGGGTGCGCTGAGAAACTGACACACGTCGGAGAAGACATCGTGAACAAAGGCGGAGACTGTGACGTGCGCGAGCTGATGGCGCGCTTTACCACGGAGTTCATCGGCGCGTGCGGCTTCGGCATACAGATGGACACCATCAACGACGAGCACTCACTCTTCAGAGATTTGGGCAGAAAAATGTTCAACAGATCGACGAAAATTCATTTCCTCGTAGCACTATATGATCTATTTCCTGAGTTTGGGACATTCTTTCAAAAAACATTCAAGGAGACATTCCTACTGGATTCCATGACAGAGATAGTTCAAAATATTCGTAAACAACGTAACTTCAAGCCCATTGGCCGAAATGACTTCATTGATTTACTTTTAGAGCTGGAAGAAAAGGGTAAAATCACAGGGGAGTCGGTGGAGAAGAGAACTCCGGATGGAAAGCCAATGCAAGTAGAAATGGAAATGGATCTGACATGCATGGTGGCGCAGGTGTTTGTGTTCTTCGCAGCAGGGTTCGAAACTTCCTCATCGGCAACGAGTTTCATGTTACATCAGTTGGCATTCCACCCAGAAGAACAAAAGAAAATTCAGGAAGATATCGATAGGGTGCTTGCGAAGTACAACAACCAACTATGCTATGACTCGATCTCTGAAATGAAAGCATTGAGTAACGGGTTCAAGGAGGCCATGAGAATGTTCCCCTCTCTCGGCACCTTACACAGAGTGTGTGCAAAGAAGTACAAGATCGAAGAGTTGGGCATCACAATCGACCCAGATGTGAAGATTATCATCCCACTGCAAGGCATACAAAATGACGAGAAATATTTTGAACATCCGGAGAAGTTCATGCCGGATAGATTCAACGATACATCAGAGGAAAGAAATAAGTTTGCATACATGCCTTTCGGTGAAGGACCTCGGCAGTGCATCGGTGAGTAGCTAAAACATGACTTATGAACTAGACACTTATTATTCTAGACCGCGTTGTTCAGTTACACCTTTCAAATTATAAAGTCTAATTTACACTTGCGAGTTAATAGCCCACTAACGCAAAATAATAATTCAAACTTATCCAATTCAGTCACAGTTTTATACGAAAGGCAAACAAATAAGAATTATATTCCTAACCTTTAATGTTTTTGTCTTCCAGGTGCTCGACTGGGTGAGATGCAGTCGCTAGCAGGACTTGCAGCAGTACTGCACAAGTTCAGCGTGGAGCCGACGACGACCACCAAGAGGTACCCAGATATCAACCACGGCAGCAACATTGTGCAATCCATCGAGGGAGGTCTGCCACTTAAACTGAAACTTAGGAGCAAGGAAAATAATGTTGCTGCATGAGGAACAACTACTATAGATCTCCATTCAAAGGAACATCATGAGCTAAAGTTACATAGGTAGATAAATGAAGATGATAATTGAACAGCACCTAATATTGAATTAGGTGTTTTTAATATTCAAGGTTATTTATTAACACCCAGTGATACCAAAATAAGCAATAACAATTACGTAGAACGAAATTGTATGGTCATTAAACTTGCTATGACCATTAATTTTATAGGGTTGAAAATAAGTATTACAGTTGTTGGTTCTATTATTCTTATAATTGTAATTTTTAAGCACCTAAAATATATAAGAAAAAAATTATAATAAGTTTCTTTTATTTGCATCTATTTTCATAGTGAGTCATCCGGACTGGTTTAGCCGACTGATAGGAATTTTAATAACTTGCAATTATCTTCGATCTTTGATATTTGAGTGTTTCTCAAGAATTAAAAATAAGTTTTATTCGTATACAAAGTTTAAAGACATCCAATCGTTATTACCTTCTGTTTAAATAAAATTAATGTTAATGTAAATCAATTCAGGGCCATGAAATATAAATTTGCTGATATCTGGCATTTCTGCCTCCTTTTGTCTTTATAATTTATAGTTCCAGTTGTTCTGAGGAAACTGTTCCCGATCCAAGATAAGTAGCAGTCTCTAACATTCAATCTGTTTCATGACAATGCATATTATCTATGCACCAAACTTCCATTCACGATTAAATAGAAAAATCACATAAGAATTCATTTGCCTACTGTTATTAGGTATTAAATACAAGCTGATCCCGCGAACTTCGTATCGTTCAAACCTTCCCTGGCTCAATCGGCCCAGCCGTTCTCGAGTTTTAATCAGACTAACGAACAACAATTCATTTTTATTTATACAGATTTAGTTTACAAATATAATATCTATCTCGGTTTCCCGTTTTACCGTTAATAAATCAACAGACTGATATGATTTAGTATCCCTAACATTAAACGTTGCAATTCACACCGTATTTATGCAGATATTCAATGAACGATAACAAGTTACAACGTTCACTACACTGTTTATTCAGTCTCGCGCGCCTGGCCGCGGCAACAACATGCTTCTCTTACTTATACCAATTGTGCTCGTGCTCTTATACTACTACACCACCAGGAACCATGACTACTGGGAGAAACGCAACATCAAGCACGAGAAACCAATACCCGTATTTGGAACGCTCTACAACAATGTATTTGCCAAGAAAAGCATCACAGAAATATCCGTGGAGCTATACCGCCAGTATCCCAATGAGAAAGTTGTGGGAATATACAGAGGAACTACACCAGAACTCATTATTCGAGATCTGGATATAGCAAGGAAGATCCTTAATGTAGACTTCGCATATTTCTACCCCAGAGGACTGGGCAGGAACCCCAAATATGAGCCCGCGTTTCTGAATATCTTCCATGTTGACGGCGACACGTGGAAATTGTTACGACAGCGTCTAACATCAGCGTTCACGACCGCCAAACTGAAAAGCATGTTCCCACTAGTAGTGGCGTGCGCTGAGAAACTGCAAAATGTGGGCGAAGATATCGTGAACAAGGGCGGAGACTGTGACGTGCGCGAGCTGATGGCGCGGTTCACCACAGAGTTCATCGGCGCGTGCGGCTTTGGTATTCAAATGGACACCATCAACAACGAGCACTCTCTCTTCAGAGATTTGGGCAGAAAAATGTTCAACAGATCGTGGAGAAATTTCGTCCTCGTACCACTCTACGATTTGTTTCCCGACTTCAGAACAGTTTTTCAAAAAATATTGCACGAGCCGTTCCTGTTGGATTCCATGGCCGAGATTGTTCAAAATATTCGCAAACAACGTAACAACAAACCAATTGGCCGAAATGACTTCATTGATTTACTTTTAGAAGTAGAAGCCAAAGGCAAGATGACAGGTGAGTCAGTGGAGAAAAGAACAGCAGATGGAAAGCCAGTGCAAGTAGAAATGGAAATGGATCTGACATGCATGGTGGCGCAGGTGTTCGTGTTCTTCGCAGCAGGTTTCGAAACTTCCTCATCGGCAACGAGTTTCATGTTACATCAGCTGGCATTCCACCCAGAAGAACAAAAGAAAATTCAGGAAGATATCGATAGGGTGCTGGCGAAGTACAACAACCAACTGTGCTATGACTCGATCTCTGAAATGAAAGCATTGAGTAACGGGTTCAAGGAGGCAATGAGAATGTTCCCTTCTCTCGGCACCTTTACACAGAATGTGTGCTAAGAAGTACAAAATCGAAGAGTTAGGCATCACAATCGATCCAGATGTGAAGATTATCATCCCAGTGGAAGGTATACAAAATGACGAAAAGTATTTCGAAAACCCCACTCAATTTAAACCGGATAGGTTCAACGATCTGTCTGAGGAGAGACATAAGTTTGCATACATGCCCTTCGGTGAAGGACCTCGGCAATGTATAGGTGAGTAGTTTTTATATTGTTTTTCTCACAAAGACCAAAGACCTTTGTTTCGTTAAGTCATTTATAGTAATCTAACCTTGATATTGAAGCTGATAACTTTCACCAAAATACTACCTTCAATAAAATGATACGATGGCCAACGCTTTCCATATTATCGGCTCTAGATTAGAGCAATAAAATTATTCATATGGAACCGAAATTTTCTGTCAAGTGGTATCCATGTGGATAACTTCAATATGATAAGACAGCAACCCAACTGCAATATTCTTGGATACCTATGCCTTTCGATATTTGAGGACCTTTTTTTGGGGCTACAATTATGATTTATTTAAATACTAATAAGGAAGGCCTTAGTCAGTAACAGTCTGCCATTTCTTTTCTATCCATCTTCAACTTTAAGGTTCTCTAAGGTTTCTTGTCAATGATTTTCTCATCAGGTGCTCGACTGGGTGAGATGCAGTCGTTAGCAGGACTTGCAGCAGTACTGCAAAAGTTCAGCGTGGAGCCAGCAGCGACCACCAAGAGGTACCCTGAGGTGAACCACGGCAGTAATACGGTGCAGTCGATCAAAGGAGGACTGCCACTGAAGTTGACTTTGAGGAATAAATAGACTTTCAAAGAACCCTCCTATTTTTTGTTCCAAAAGCCATGATGTCCACCAATGCTCTATTTGCAAAATAAACCACTTTACTATCCACGAAGCAGACACAAGAGTCAAACAATTGGTGTGTATCAAATATGATTCATGTGACAGGGAAAATTATTAAGAGCTCACCATTTTTTGGAACCCGCAGAAGAAATCAAATCCTACGAATTCATCCAGACTTGAATAGAAGAGTTCTATTGAAAATAGTAAAGGCTCCGAAAACCATATAACCATATAAAAGGGTTAAAGCCCTTAATCAAGAGCTCTTAATCGTACCTAAATAGATAATTATTATCGTAAAGTAAATCAATAAATACTAAGATAA

>Amino acid sequence of HarmCYP6AN1 in Fig. 2

MIFLLLLPIALVLLYKYTTRNHDYWEKRNIKHEKPIPVFGTLYDNVFAKKSIIEVAADLYRQYPNEKVVGMFRGNTPELVIRDLEIVRKILNVDFAYFYPRGIGRNPTYEPLFMNLFHAEGDSWKLLRQRLTSTFTTAKLKKMFPLVVGCAEKLTHVGEDIVNKGGDCDVRELMARFTTEFIGACGFGIQMDTINDEHSLFRDLGRKMFNRSTKIHFLVALYDLFPEFGTFFQKTLKETFLLDSMTEIVQNIRKQRNFKPIGRNDFIDLLLELEEKGKITGESVEKRTADGKPVQVEMEMDLTCMVAQVFVFFAAGFETSSSATSFMLHQLAFHPEEQKKIQEDIDRVLAKYNNQLCYDSISEMKALSNGFKEAMRMFPSLGTLHRVCAKKYKIEELGITIDPDVKIIIPLQGIQNDEKYFEHPEKFMPDRFNDTSEERNKFAYMPFGEGPRQCIGARLGEMQSLAGLAAVLHKFSVEPTTTTKRYPDINHGSNIVQSIEGGLPLKLKLRSKENNVAA

> Amino acid sequence of HassCYP6AN1 in Fig. 2

MIFLLLLPIALVLLYYYTTRNHDYWEKRNVKHQKPIPVFGTLYDNVFAKKSIIEMAADLYRQYPNEKVVGMFRGNTPELVIRDLEIVRKILNVDFAYFYPRGIGRNPTYEPLFMNLFHAEGDSWKLLRQRLTSTFTTAKLKKMFPLVVGCAEKLTHVGEDIVNKGGDCDVRELMARFTTEFIGACGFGIQMDTINDEHSLFRHLGRKMFNRSTKIHILVALYDLFPELGTFFQKTLRETFLLDSMTEIVQNIRKQRNFKPIGRNDFIDLLLELEGKGKITGESVEKRTADGKPVQVEMEMDLTCMVAQVFVFFAAGFETSSSATSFMLHQLAFHPEEQKKIQEDIDRVLAKYNNQLCYDSISEMKALSNGFKEAMRMFPSLGTLHRVCAKKYKIEELGITIDPDVKIIIPLQGIQNDGKYFEHPEKFLPDRFNDASEERNKFAYMPFGEGPRQCIGARLGEMQSLAGLAAVLHKFSVEPTTTTKRYPDINHASNIVQSIEGGLPLKLKLRSKENNVAA

>5' flanking sequence of *HarmCYP6AN1* (-114 to +763) in Fig. S1

ATAGTCTGACTTTTGTCATAATACTCTAAAATTTTTGTCTACGAGCGACCGAACAAAGTCCAAAGTCTATATTGGACCGAGATATATAATACTGAGTTATGAGGCTCTCAACTCAAACTGGTTTGAACCATCCAGCTAGAACAACCTATGCTAGAATTTCGCGAGTTTTTGCCAAATAGTGATCGAAATAAACATTTTTTAACACGGAAGAACTGAGGTATGTATTGTTTGATAAAGTGTTTTCTTAACCTTTGGTAGTGTAAATAAGTTATTATCAAGTGTAAGGAGATTGTGTTAAAAATAGACAACTTGAGTCCAAAGGTCGATTTTTAGCTTTATTTGGTGTCTGAAGAAAGTAACTTTCTTTAGTGAACGGGTAAAGTGGCATTTATGATCAAAGTCTAGATCAATCATGATAGTATCATGATTTATTTGTGTTTATTAACTTATCGGTGTTTTATTGGGTGTTAAGCCTTTGTGTATCGTATCAAAACATGTTCAAAATCTCATCTATTTTTGAAGTAGGTAATTATTTGCTACATCATTTACATTTTCCGATTTTTTTTTGCAAATTATGCTAAACTTTATTTTTATTGTATCGTGGGTACCTACCTTCATTGTAATATCTTATTCTTGTCTCACCTTTGATAACTATTTACTATTTTATTACTAAGGAAGGTGATAAGGACATGTTCTTATCACCTTGTCTAGGCATACAGCTGGTAGAATATTGAGATAATAGTTGAAAGATAACAACGTGTAAAAATATATTCCAGTATCAACCAGTTACACAACTTGTGTGAGTAAACAGCACTGTTGAAATTCAATACAATTATCGTATCTTTTTGTAGTTAATCGATAGTTTCTTCAATGCCC

>5' flanking sequence of *HassCYP6AN1* (-114 to +784) in Fig. S1

ATAGTCTGACATTTGTCATATTACTCTAAAATTTTCGTCTACGAGCGACCGAACAAAGTCCAAAGTCTATAATGTACCGAGATATATAATACTGAGTTGTGAGGTTCTCAACTCAAACTGGTTTGAACCATCCAGCTAGAACAACCTATATTTTAAAATTTCGCGAGTTTTTGCCAAATAGTGATCAAAATAAACATTTTTTTAATACGAAGAACTGAGGTATGTATTATTAGATAAAGTGTTTTCTTAACCTTTGATAGTGTAAATAAGTTATTATCAAGCGTAAGGAGATTGTGTTAAAAATAGACAACTTGAGTCCAAGGGTCGATTTTTAGCTATATTTGGTGTGTGAAGAAAGTAAATTTCTTTAGTGAACGGTTAAAGTGACATTTATGACCAAAGTCTAGATCAATCATGATAGTATCTTGATTTATTAGTGTTTATTATCAACTTATCGGTGTTTTATTGGGTGTTAAGCCTTTGTTTATCGTATCAAAACATGTTCAAAATCTCGTCTATTTATGAAGTAGTTAATTTACTTGCTACATCATTTACATTTCCCGACAATTTATGCAAATTAAGCTAAACTTTATTTTTATTGTATCGTGGGTACCTACATTGTAATATCTTATTCTTGTCTCACCTTTGATAACTTTTTACTTTTAGGTATATTACTAAAATATAAAGACAATGTGATAAGAAAATGTCCTTATCACCTTGTCTAGATATTTTAAGTACAGCTAGTGGAATATTGAGATAATAGCTGAAAGATAACAACGTACTTATAAAAATATACTCCAGTATCAAGCACTTACACAATTGTGAGTAAACGGCACTGTTGAAATTCAACATAATTATCGTACCTTTTTGTAGTTTATCGATAGTTTCTTCAATGCCC
